# Metabolic Dynamics in *Escherichia coli*-based Cell-Free Systems

**DOI:** 10.1101/2021.05.16.444339

**Authors:** April M. Miguez, Yan Zhang, Fernanda Piorino, Mark P. Styczynski

## Abstract

The field of metabolic engineering has yielded remarkable accomplishments in using cells to produce valuable molecules, and cell-free expression (CFE) systems have the potential to push the field even further. However, CFE systems still face some outstanding challenges, including endogenous metabolic activity that is poorly understood yet has a significant impact on CFE productivity. Here, we use metabolomics to characterize the temporal metabolic changes in CFE systems and their constituent components, including significant metabolic activity in central carbon and amino acid metabolism. We find that while changing the reaction starting state via lysate pre-incubation impacts protein production, it has a comparatively small impact on metabolic state. We also demonstrate that changes to lysate preparation have a larger effect on protein yield and temporal metabolic profiles, though general metabolic trends are conserved. Finally, while we improve protein production through targeted supplementation of metabolic enzymes, we show that the endogenous metabolic activity is fairly resilient to these enzymatic perturbations. Overall, this work highlights the robust nature of CFE reaction metabolism as well as the importance of understanding the complex interdependence of metabolites and proteins in CFE systems to guide optimization efforts.

Abstract Figure

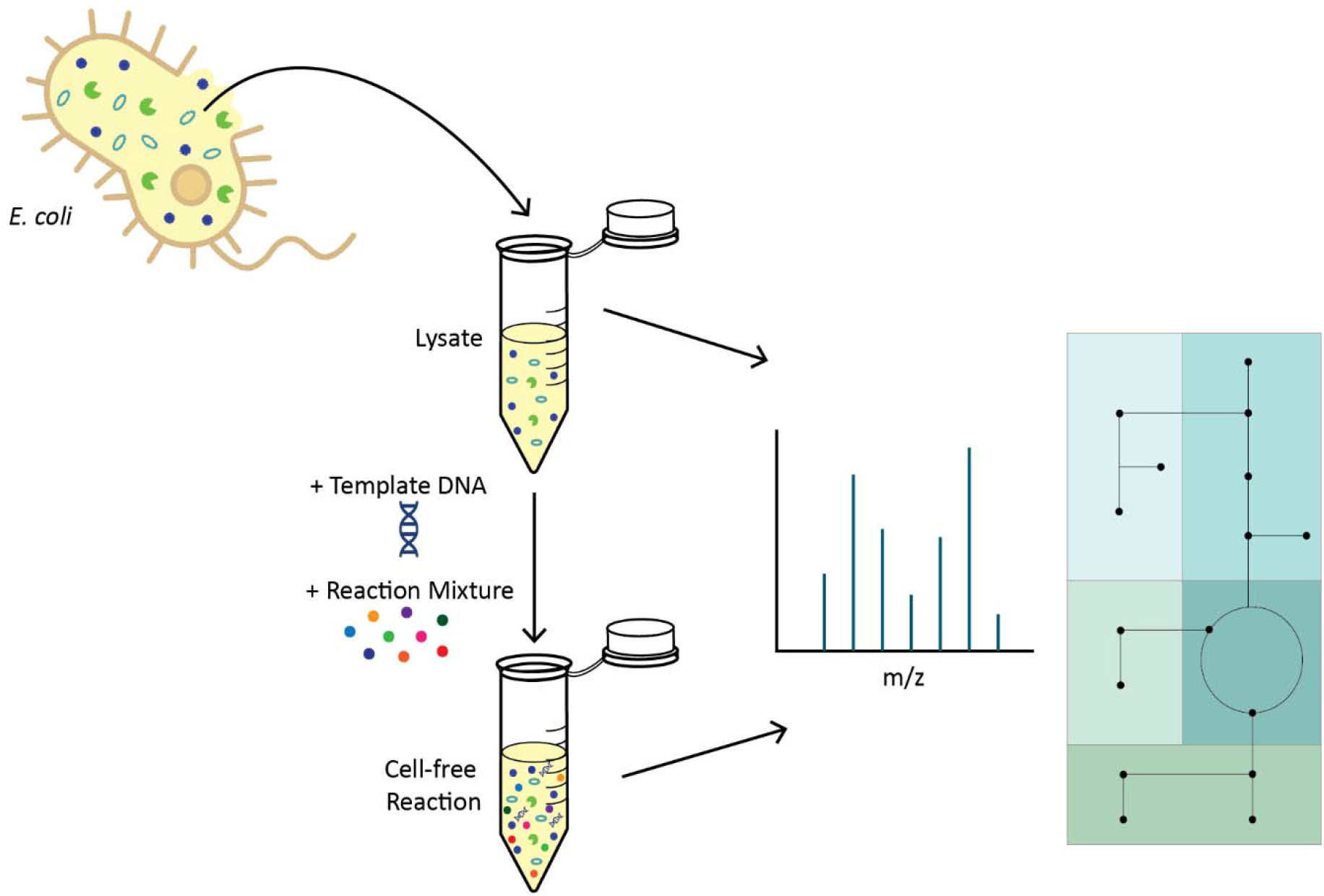

## INTRODUCTION

Advances in metabolic engineering (the engineering of cells to produce valuable chemicals) have been driven in part by a deeper understanding of cellular metabolism, the energy-supplying chemical reactions within living organisms. This deeper understanding has enabled identification of key (sometimes distal) areas of metabolism that inhibit or enable production of target molecules. Cells have been used for the production of biotechnologically-relevant products such as lactic acid, 2,3-butanediol, and isoprene at industrial-scale titers^1^, and have even successfully been used for the production of molecules at specific concentrations or times through the use of precision metabolic engineering^2, 3^ or dynamic metabolic engineering^4, 5^, respectively. However, the use of whole cells as biocatalysts has some inherent limitations, as the natural programming of these living organisms leads to some fraction of substrates, energy, or cofactors being funneled to support the cells’ growth and survival rather than the design goals of an engineer.

A promising alternative approach to producing chemicals without the metabolic burdens of cellular maintenance is the cell-free expression (CFE) system. CFE entails the use of *in vitro* systems containing transcription and translation machinery, either from purified recombinant elements (PURE) or derived from the crude protein lysate (extract) of whole cells. These systems can be easily engineered for biocatalysis and biosynthesis. Because the native genetic programming (genomic DNA) is removed from CFE systems, typical energy- and metabolite-intensive cellular requirements like cell replication are avoided. Moreover, the lack of cellular membranes removes transport limitations than can hinder biocatalysis and allows for continuous monitoring and manipulation. These advantages have led to the successful implementation of CFE systems for the production of small molecules and proteins in broad application areas ranging from therapeutics and biosensors to biofuels^6–9^.

One of the most widely used CFE platforms is the *Escherichia coli* lysate-based system. The cellular lysate provides the major components that allow for transcription and translation from an added DNA template that codes for an enzyme, pathway, or genetic circuit. The lysate is supplemented with a reaction mixture that provides additional biochemical building blocks and energy-providing metabolites as well as cofactors and salts. Both the lysate and the reaction mixture have been the focus of many significant optimization efforts^10–13^.

However, reproducibility, scalability, and standardization challenges still exist due to the complex and poorly-understood impacts of the preparation and constituents of these reaction components on CFE system performance. Crude cell lysate is a complex mixture, and it is unknown how much its endogenous enzymes and metabolites impact the productivity of a CFE reaction. One prominent hypothesis is that CFE reaction efficiency and lifetime is significantly (or even predominantly) affected by the depletion or accumulation of specific metabolites that affect protein synthesis^14, 15^, though the identity of the molecules involved is not known. While it is known that many central carbon metabolism enzymes are present in *E. coli*-based cell-free lysates^16–18^, it is unclear how they impact metabolic processes downstream of central carbon metabolism and alter the transcriptional and translational capacity of a CFE reaction **(Figure 1)**. Unfortunately, unlike in living whole-cell biological systems, we have minimal understanding of the endogenous metabolism in CFE systems, prompting a serious need for characterization of their metabolic dynamics.

**Figure 1.**
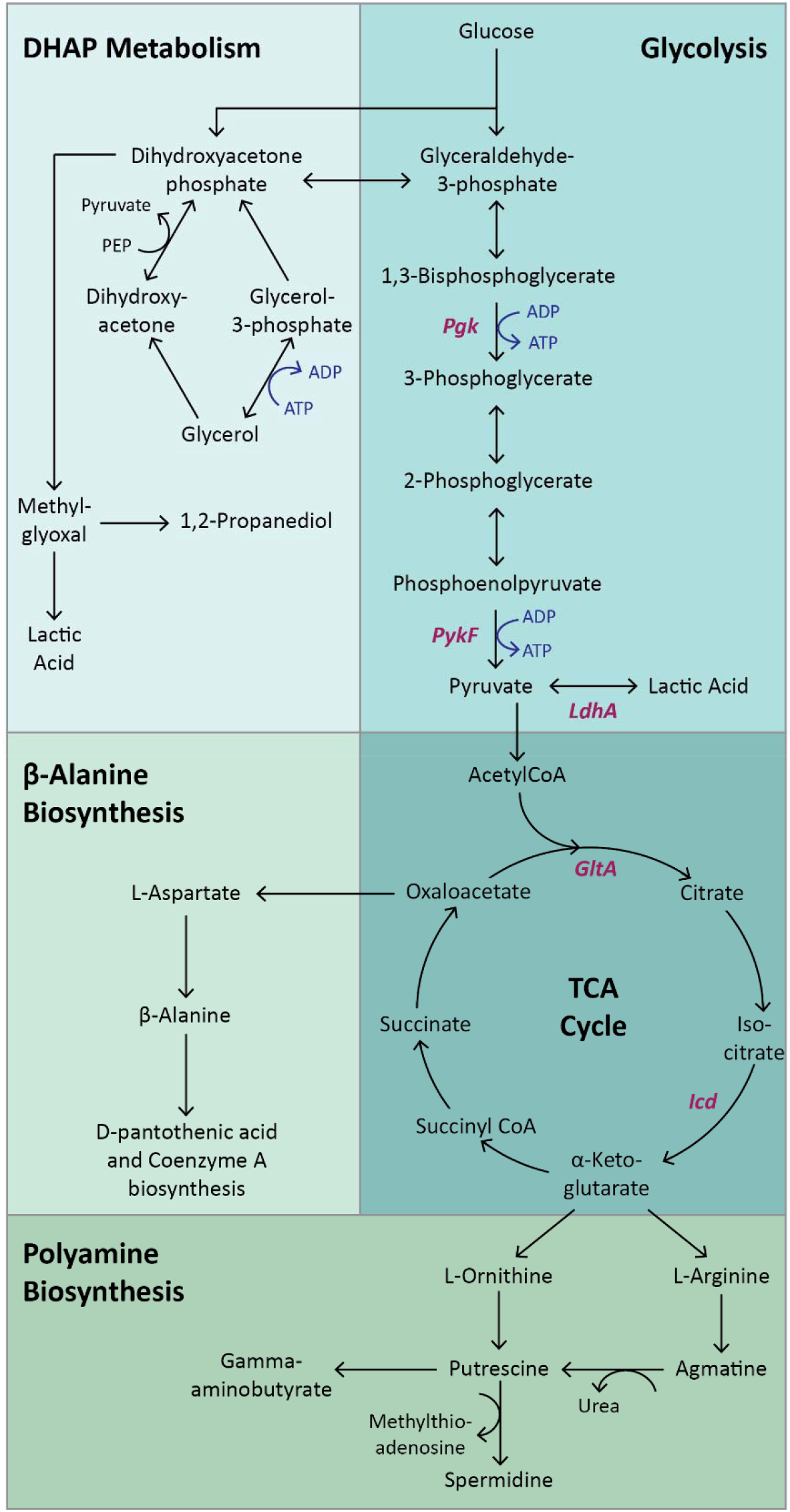
An overview of metabolic pathways relevant to this work; enzymes of interest specifically discussed in this paper are noted in italics.

Systems biology and the associated “omics” tools that have matured tremendously in the past two decades provide the potential for broad-ranging insight into such problems at a cell-wide scale. Metabolomics—the study of the small molecule intermediates of the chemical reactions within a living organism—is a particularly useful approach to measure the temporal metabolic changes in whole cells and CFE systems alike^19, 20^. Hundreds to thousands of small molecules can be tracked and quantified using analytical approaches such as mass spectrometry and NMR, with downstream informatics used for analysis and biological interpretation.

In recent work, we used metabolomics to assess the impacts of differently prepared *E. coli*-based CFE lysates and found that the reactions made from these different lysates were metabolically distinct^21^. To our surprise, we found that the metabolite-level changes at the end of a CFE reaction due to endogenous metabolic activity in the reactions eclipsed any metabolic changes due to protein synthesis. These findings highlight the complexity of CFE systems, our lack of understanding of their metabolic underpinnings, and the resultant need for broader metabolic investigation of CFE systems. Deeper understanding of CFE metabolism would facilitate not only optimization efforts, but also rational approaches to resolve reproducibility, scalability, and standardization issues in CFE systems.

Here, we used metabolomics to more broadly characterize the metabolic profiles of *E. coli*-based CFE systems. In particular, we aimed for a deeper characterization of the metabolic dynamics of these systems, as previous analysis focused on long end-point times and thus may have missed critical dynamics during protein synthesis and as CFE activity declined. We also deconstructed the CFE reaction into its constituent components for metabolomic analysis, in an effort to more clearly pinpoint the source of metabolic changes observed in complete CFE reactions. We explored the effects of lysate pre-incubation and how changes to the sonication energy input during lysate preparation affect both endogenous metabolism and protein synthesis in CFE reactions. Finally, we used the information from these studies to select native metabolic enzymes to supplement in CFE reactions in an effort to alter metabolic activity and thus protein yield of a reaction.

## RESULTS/DISCUSSION

### Metabolic profiles change throughout CFE reactions

Our first goal was to characterize the metabolic dynamics in a CFE reaction for the duration of protein synthesis, as previous studies had focused on reaction endpoints. We prepared the lysate from exponentially growing BL21 cells in 2×YTP media; the cells were lysed via sonication, and the lysate was post-processed with a run-off reaction and dialysis (see Methods for details). A CFE reaction was assembled comprising this lysate, a small-molecule reaction mixture, and the CFE plasmid pJL1s70 to drive expression of green fluorescent protein (GFP) from an *E. coli* σ^70^ promoter. GFP production was measured and samples were collected for metabolomics analysis at 0, 0.5, 1, 2, 4 and 6 hours. Metabolomics samples were prepared by precipitating proteins and analyzing the remaining metabolite mixture using two-dimensional gas chromatography coupled to mass spectrometry (GC×GC-MS) after sample derivatization. The resulting instrument output was processed with a computational workflow resulting in relative abundances for 276 putatively identified and unannotated metabolites that were used for downstream analysis.

For a systems-scale analysis of the temporal metabolic changes in the reaction, we analyzed the resulting data with principal component analysis (PCA). PCA is an unsupervised dimensional reduction method that identifies the modes of the data that capture the most variance, known as the principal components (PCs). Generally, separation of sample groups in the first few principal components indicates prominent differences in metabolite profiles, which can include metabolites with individually significant differences between groups as well as metabolites with individually insignificant but correlated changes between groups. For the time-course CFE profiles, the majority of sample group separation is captured in the first principal component (PC1), reflecting a monotonic change in metabolic state over the course of the reaction (**Figure 2B)**. The fact that the 0 hour and 0.5 hour samples are the most separated consecutive timepoints in PC1 suggests that a large portion of the metabolic changes likely occurred in the first half hour of the reaction; however, the separation of later timepoint samples in this same principal component suggests that metabolite levels continue to change throughout the entirety of the reaction. Surprisingly, metabolite profiles at 4 hours separate from those at 6 hours in both PC1 and PC2, showcasing that significant metabolic activity continues even as protein synthesis is concluding **(Figure 2A)**.

**Figure 2.**
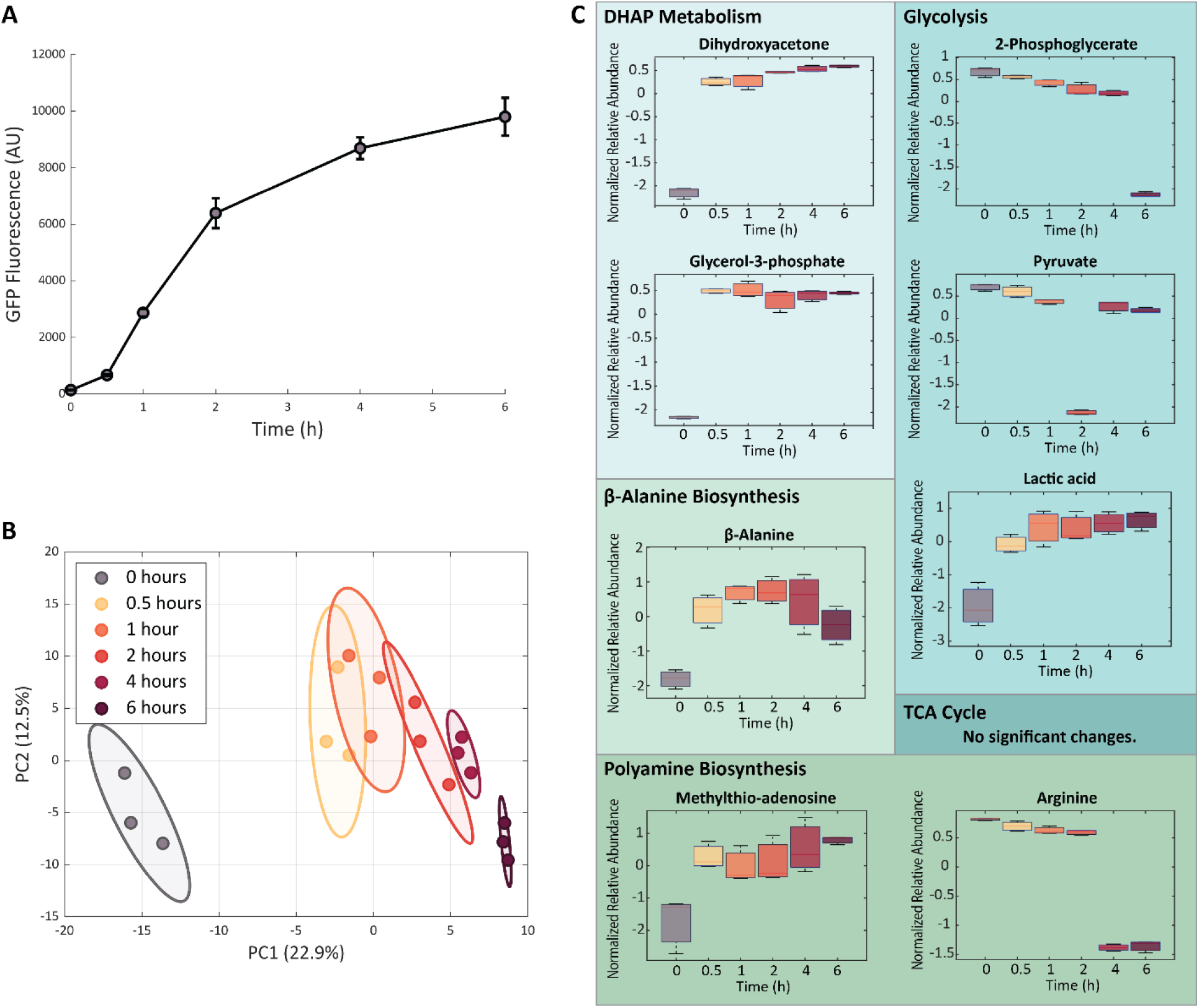
Temporal profiles of CFE reactions during protein production. (A) GFP production (measured via fluorescence) slows down at around 4 hours. Error bars represent standard deviation of triplicate reactions. (B) Samples from different timepoints separate from one another, with PC1 values increasing with reaction time. Colored ellipses represented 95% confidence intervals for each group, and the plotted samples are triplicate reactions. (C) Profiles of metabolites involved in glycolysis, dihydroxyacetone phosphate (DHAP) metabolism, β-alanine biosynthesis, and polyamine biosynthesis pathways change over the course of a CFE reaction. Box and whisker plots depict the normalized peak areas, which are transformed using a generalized logarithm (base 2) and autoscaled. Red lines are the medians, boxes span the second and third quartiles of values. Error bars represent standard deviation of triplicate reactions.

### Levels of multiple key metabolic pathways evolve over the course of a CFE reaction

We then identified individual metabolites with significant changes during the reaction using one-way analysis of variance (ANOVA). We found major metabolic changes in central carbon and amino acid metabolism. Specifically, we detected significant changes (using False Discovery Rate (FDR)-corrected *p*-values < 0.05) to metabolites involved in glycolysis, dihydroxyacetone phosphate (DHAP) metabolism, β-alanine biosynthesis, and polyamine precursor biosynthesis **(Figure 2C**). There were significant decreases in the abundances of the glycolytic intermediates 2-phosphoglycerate (2-PG) and pyruvate and a continuous increase in the fermentation product lactic acid, potentially from the conversion of pyruvate **(Figure 1)**. These observations were unsurprising, as glycolysis is the primary pathway for CFE reactions to create ATP, and it has previously been shown that most glycolytic enzymes are present in *E. coli*-derived lysates^16–18^.

Changes in other glycolytic byproducts involved in DHAP metabolism were more unexpected. DHAP is a glycolytic intermediate that has various routes for conversion: (1) isomerization into glyceraldhyde-3-phosphate to enter glycolysis, (2) conversion into the glycerol-associated metabolites dihydroxyacetone or glycerol-3-phosphate, or (3) breakdown into the highly toxic molecule methylglyoxal that can in turn become the less toxic metabolites 1,2-propanediol or lactic acid^22^. Two metabolites within this pathway, dihydroxyacetone and glycerol-3-phosphate, accumulated significantly during the CFE reaction. These metabolites are substrates for or products of glycerol, but we did not detect any significant changes to glycerol **(Figure S1A)**. The increase in lactic acid could potentially result from conversion of DHAP to lactic acid due to accumulation of methylglyoxal. Methylglyoxal is known to be highly toxic to cells due to its ability to interact with DNA^23^, though it was not identified in this data set. If present in the CFE reaction at appreciable levels, methylglyoxal could interact with template DNA and inhibit expression.

Beyond this more central portion of carbon metabolism, some sections of amino acid metabolism also exhibited significant temporal profiles during the CFE reaction, including β-alanine biosynthesis. β-alanine is the product of L-aspartate, which is synthesized from oxaloacetate (OAA) in the tricarboxylic acid (TCA) cycle. Although L-aspartate did not significantly change **(Figure S1B)**, we found that β-alanine increased at the beginning of the reaction and remained relatively constant after the first hour. This metabolite and its interesting dynamics are notable for a number of reasons. First, it is the precursor to pantothenic acid (vitamin B_5_) and thus to Coenzyme A, an essential cofactor for many key metabolic pathways including the TCA cycle, fatty acid biosynthesis, and acetyl-CoA production^24, 25^. Second, and perhaps even more noteworthy, we have previously shown that supplementing β-alanine to a CFE reaction increases protein expression^21^, indicating that its levels are (either directly or indirectly) important to CFE.

Polyamine biosynthesis was another section of amino acid metabolism with significant temporal profiles. The polyamines putrescine and spermidine are known to be extremely important for processes in living cells due to their key roles in cell-to-cell signaling, cell division, cell motility, and synthesis of DNA and proteins^26^. They are also components of the CFE reaction mixture. Polyamine biosynthesis begins with the molecules L-ornithine or L-arginine, which both can be derived from α-ketoglutarate in the TCA cycle; byproducts of the pathway include urea and methylthio-adenosine. Although ornithine and putrescine remained constant in our measurements **(Figure S1C-D)** and spermidine was not identified, the precursor metabolites L-arginine and methylthio-adenosine decreased and increased, respectively. Interestingly, the two TCA metabolites we detected (fumarate and malate) did not change significantly **(Figure S1E-F)**, potentially indicating that TCA molecules are not significant precursors for β-alanine or polyamine biosynthesis, likely due to the excess of amino acids in the reaction mixture.

### The metabolic profile of the lysate alone changes with time

To begin to pinpoint the source of the metabolic changes that occur in complete CFE reactions, we sought to separately identify metabolic changes in each of the two main constituents of the reaction: the lysate and the reaction mixture. To that end, we measured metabolite profiles in the lysate and in the reaction mixture when separately incubated under otherwise normal reaction conditions. Water was added to the lysate and reaction mixture samples to bring them to the same volume as a CFE reaction, and both were incubated at 37 °C and collected for metabolomics analysis at 0, 1 and 4 h. Data processing yielded 247 and 303 known and unannotated analytes for the lysate and reaction mixture samples, respectively, to be used in further analyses.

Principal component analysis revealed no separation of reaction mixture metabolite profiles at the different timepoints but did yield distinct separation of lysate samples in PC1 **(Figure 3A and B**), consistent with our expectations for both sample types. We expected the small molecules in the reaction mixture to be stable without any enzymes present, and only twelve metabolites (**Figure S2**) were identified as significantly changing using ANOVA (most of which were not annotated and were likely a result of poor chromatographic peak resolution). We also expected the enzymes in the lysate to cause changes in metabolite profiles. However, it is worth noting that the lysate had already undergone a “run-off reaction” (to degrade host RNA/DNA) and dialysis^14^, making the presence of a lysate metabolome and its potential for significant transformation somewhat surprising.

**Figure 3.**
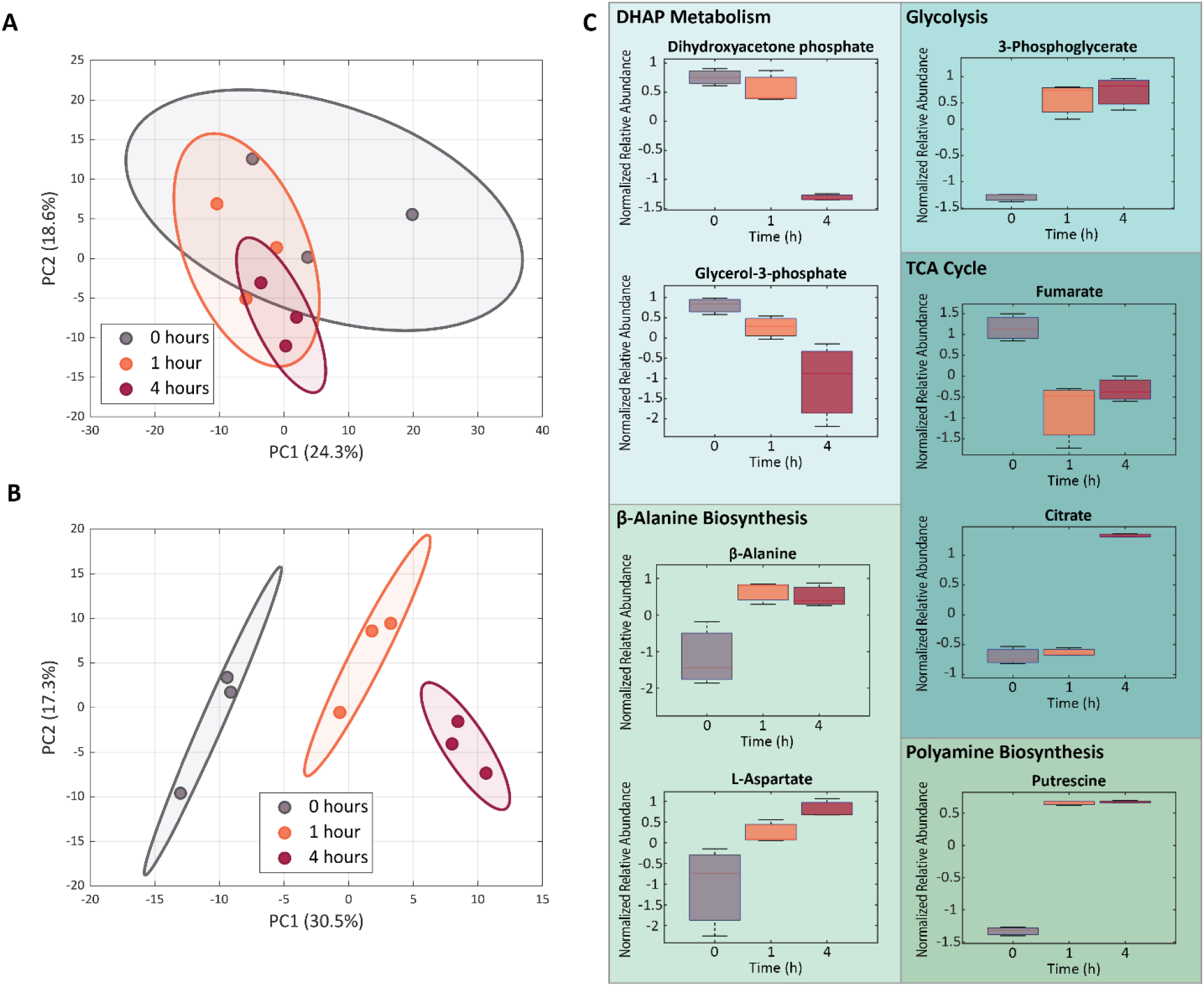
Metabolic changes in incubated reaction mixtures and lysates. (A) PCA plot for the incubated reaction mixture samples, showing no clustering of or separation between timepoints. (B) PCA plot for the incubated lysate samples, showing distinct separation between metabolite profiles at each time point. For (A) and (B), colored ellipses represented 95% confidence intervals for each group, and the plotted samples are replicate reactions. (C) Metabolites involved in glycolysis, DHAP metabolism, the tricarboxylic acid (TCA) cycle, β-alanine biosynthesis, and polyamine biosynthesis pathways change during lysate incubation. Box and whisker plots depict the normalized peak areas, which are transformed using a generalized logarithm (base 2) and autoscaled. Red lines are the medians, boxes span the second and third quartiles of values. Error bars represent standard deviation of triplicate reactions.

The metabolic pathways with significant changes in the lysate were similar to those in the complete CFE reaction and included glycolysis, DHAP metabolism, β-alanine biosynthesis, and polyamine precursor biosynthesis **(Figure 3C)**. Additionally, molecules in the TCA cycle significantly changed.

Notably, though, many of the metabolites identified as significantly changing in the lysate alone did not have the same temporal trends as in the complete CFE reaction. Glycerol-3-phosphate, citrate, fumarate and putrescine do not trend the same way as in the complete reaction; however, the decreasing trends for glycerol-3-phosphate were not statistically significant. β-alanine does trend similarly to the complete CFE reaction, but the changes were not statistically significant. Additionally, pyruvate, lactic acid, glycerol, and urea levels did not significantly change over time **(Figure S3A-D)**.

Thus, the changes in metabolite profiles during CFE reactions do appear to be attributable to the endogenous metabolic activity of the lysate rather than chemical degradation of the reaction mixture. In fact, similar metabolites are affected in both the lysate and the complete CFE reaction, suggesting the prominent roles of similar enzymes. However, the different trends in those metabolite profiles between the two cases indicate that the surplus of molecules provided in the reaction mixture fundamentally alters the qualitative impacts of endogenous metabolic activity for those enzymes.

### Lysate incubation affects protein yield with minor impacts on CFE reaction metabolic state

We next sought to characterize the relationship between the lysate’s initial metabolic state and the productivity and final metabolic state of a CFE reaction. Previously, we demonstrated that pre-incubating the lysate with reaction mixture at 37 °C for 8 h prior to DNA template addition substantially decreased protein ouput^21^, suggesting that the endogenous metabolism of the system may affect its productivity. Since we now know (**Figures 2 and 3**) that the metabolic trends of the lysate’s endogenous enzymes can be quite different depending on whether the reaction mixture metabolites are present, this led to the question of whether pre-incubation of lysate without the reaction mixture would affect the productivity or final metabolic state of a CFE reaction.

Accordingly, we incubated the lysate without any reaction mixture for six hours at 4, 25, or 37 °C and then used these pre-incubated lysates in a CFE reaction producing GFP from pJL1s70. We selected the incubation time of 6 hours to be consistent with the timescale of protein production (**Figure 2**), allowing sufficient time for endogenous metabolic activity to approach completion. We found in small-volume microwell plate experiments that lysate pre-incubation at 25 or 37 °C, but not at 4 °C, resulted in a substantial reduction in GFP production **(Figure S4)**. We then used larger-volume reactions for fluorescence and metabolomics analysis at 0, 1 and 4 hours after the start of the reaction with pre-incubated lysate. After data processing, these metabolomics measurements yielded 424 known and unannotated analytes used in further analyses.

As seen in **Figure 4A**, CFE reactions in the larger-volume format also produce significantly less GFP when lysates are pre-incubated at 25 or 37 °C compared to at 4 °C. Multivariate analysis with PCA indicated differences in metabolic profiles between the three pre-incubation temperatures before the reaction started but a convergence of the overall metabolic profiles by the final timepoint **(Figure 4B)**, a contrast to their divergence in protein productivity. With univariate analyses, we found that most metabolic dynamics in the reactions with pre-incubated lysates were the same as in reactions with fresh lysate **(Figure 4C)**. For example, temporal trends in glycolysis and TCA cycle metabolites were independent of lysate pre-incubation temperature and identical to the fresh CFE reaction, even though levels of some of these molecules had changed during lysate incubation.

**Figure 4.**
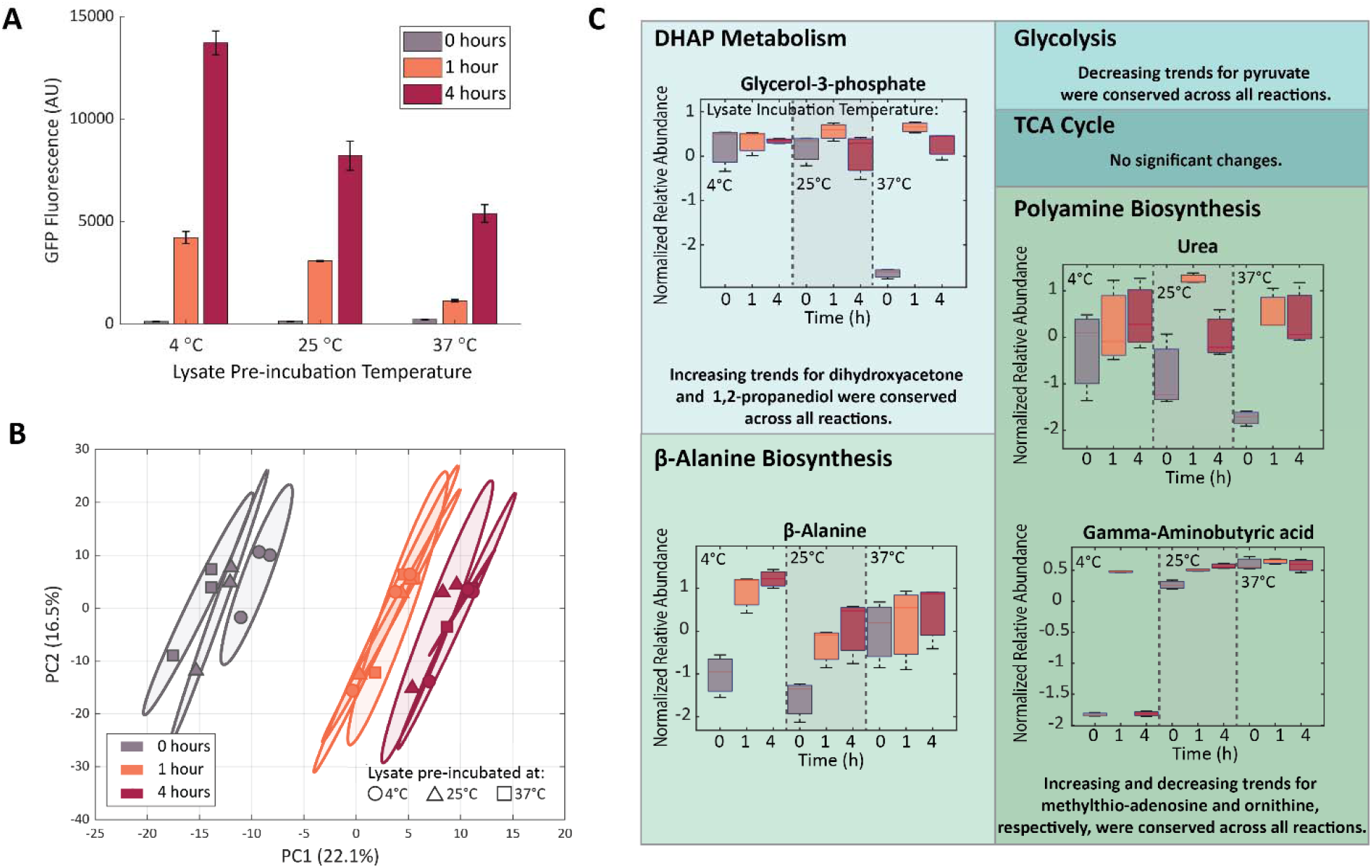
GFP production and metabolic changes in CFE reactions using lysates pre-incubated for 6 h at 4, 25, or 37 °C. (A) Increasing lysate pre-incubation temperature results in decreased GFP output. Error bars represent standard deviation of triplicate reactions. (B) Principal component analysis of metabolite profiles collected at different reaction timepoints. The samples collected at the start of the reaction separate based on lysate pre-incubation temperature, but the metabolic profiles converge as the reaction progresses. Colored ellipses represented 95% confidence intervals for each group, and the plotted samples are replicate reactions. (C) Only a few metabolites involved in DHAP metabolism, β-alanine biosynthesis, and polyamine biosynthesis pathways had different profiles for different lysate pre-incubation temperatures. Box and whisker plots depict the normalized peak areas, which are transformed using a generalized logarithm (base 2) and autoscaled. Red lines are the medians, boxes span the second and third quartiles of values. Error bars represent standard deviation of triplicate reactions. Metabolites that changed significantly with time but consistently across sample groups are noted with text.

However, a few molecules in DHAP metabolism, β-alanine biosynthesis, and polyamine precursor biosynthesis had different profiles for different lysate pre-incubation temperatures. Glycerol-3-phosphate levels stayed relatively constant when lysates pre-incubated at 4 and 25 °C were used, whereas with 37 °C pre-incubated lysates this molecule accumulated, indicating it is produced during the CFE reaction despite its downward trend during high-temperature pre-incubation. β-alanine levels increased over time for reactions with 4 and 25 °C pre-incubated lysates (similar to a fresh CFE reaction), but with 37 °C pre-incubated lysates the elevated initial concentration of β-alanine does not appreciably increase, suggesting that β-alanine biosynthesis is largely completed after 37 °C pre-incubation. Urea and gamma-aminobutyric acid (GABA, a putrescine degradation product) were also affected by increasing pre-incubation temperatures, with more accumulation at higher and lower temperatures, respectively. (Putrescine was detected but did not significantly change across samples, potentially due its abundance in the reaction mixture **(Figure S5)**.)

We also performed univariate analysis via two-way ANOVA (ANOVA2) to quantify the impact of lysate pre-incubation temperature on the metabolite profiles. We found that reaction time is the most common significant effect across the measured metabolites, though (consistent with the preceding discussion) there are also significant interaction effects **(Figure S6A)**.

Although pre-incubating lysates at different temperatures significantly affects CFE reaction protein production, the changes in metabolic dynamics caused by the use of these different lysates are small compared to the overall metabolic changes during the course of a reaction. While it was perhaps surprising that the final metabolic states of these systems with such different expression levels seem so similar, the small metabolic changes may very well still be a root cause of the changes in expression, especially since some of these molecules, such as polyamines and β-alanine, are known to have key roles in protein production. This hypothesis is supported by multiple pieces of evidence from the literature. First, the loss of protein expression with pre-incubation seems likely to be either directly or indirectly metabolic in its cause: expression in CFE reactions can be maintained for 16 to 24 hours by use of dialysis reactors^15, 27^, suggesting that expression machinery is not likely degrading during the pre-incubation. Furthermore, we previously demonstrated that supplementation of β-alanine, putrescine, and spermidine into CFE reactions can drastically alter protein production, highlighting the impact of metabolite levels on protein output^21^. Nonetheless, proteins that are temperature-sensitive or have short half-lives could be disproportionately affected by pre-incubation, which would not be detected here as enzymatic activity was not directly measured in our experiments. Additionally, the assessment of there only being “small” temperature-related changes in metabolite profiles could be biased by the finite metabolome coverage of GC-MS.

### Sonication Energy Input Significantly Impacts Protein Production and Metabolic Profiles

We next sought to further explore how susceptible endogenous lysate metabolism is to changes in its initial metabolic state via alterations to lysate preparation. Our previous studies of CFE metabolism assessed the impact of multiple steps in lysate preparation, including growth of the starter culture with or without glucose and dialysis of the lysate^21^. However, we had yet to explore one variable known to have a significant impact on lysate productivity: the sonication energy input to lyse the cells.

While sonication energy input should not impact the initial metabolite profile in the lysate, its impact on expression may be correlated with or mediated by metabolic changes, so we sought to characterize endogenous metabolism in lysates made with different sonication energies. While the exact number varies between operators due to differences in technique, a typical sonication energy input for cell lysis in our group is 250-300 J; lower input energies can be used, but may reduce lysis efficacy due to an increase in intact cells and a lower total *E. coli* protein concentration in the crude lysate^28^. We prepared lysates using sonication energies of 25, 100, and 300 J and used them for assembly of CFE reactions with the reaction mixture and the template DNA pJL1s70. Samples for fluorescence measurements and metabolomics analysis were collected at 0, 1, 4, and 12 h. Processed metabolomics measurements yielded 351 annotated and unannotated analytes for further analysis.

Different sonication energies did in fact lead to different protein yields and metabolite profiles. An energy input of 100 J unexpectedly resulted in a higher protein yield than either 25 J or 300 J at 1 h and 4 h **(Figure 5A)**. While the 25J condition had lower expression than the 300 J condition at 1 h, the 25 J condition surpassed the 300 J condition at 4 h and made comparable amounts of protein to the 100 J condition by 12 h. This temporal difference in expression profiles was reproducibly observed, but given the known operator-specific aspects of sonication energy protocol optimization, we refrain from interpreting too much from the quantitative details and instead focus on the three conditions as generally indicative of different degrees of lysis and different expression efficiency. The stark differences in protein yield reflected in those measurements were evident in our metabolomics data when analyzed using PCA **(Figure 5B)**, with significant separation of the conditions at almost all timepoints, indicating that different sonication energy inputs lead to fundamentally different endogenous CFE metabolic dynamics. Notably, the 100 and 300 J reaction samples are closer to each other in PCA space than the 25 J reactions at each time point, suggesting greater similarities in their metabolic profiles.

**Figure 5.**
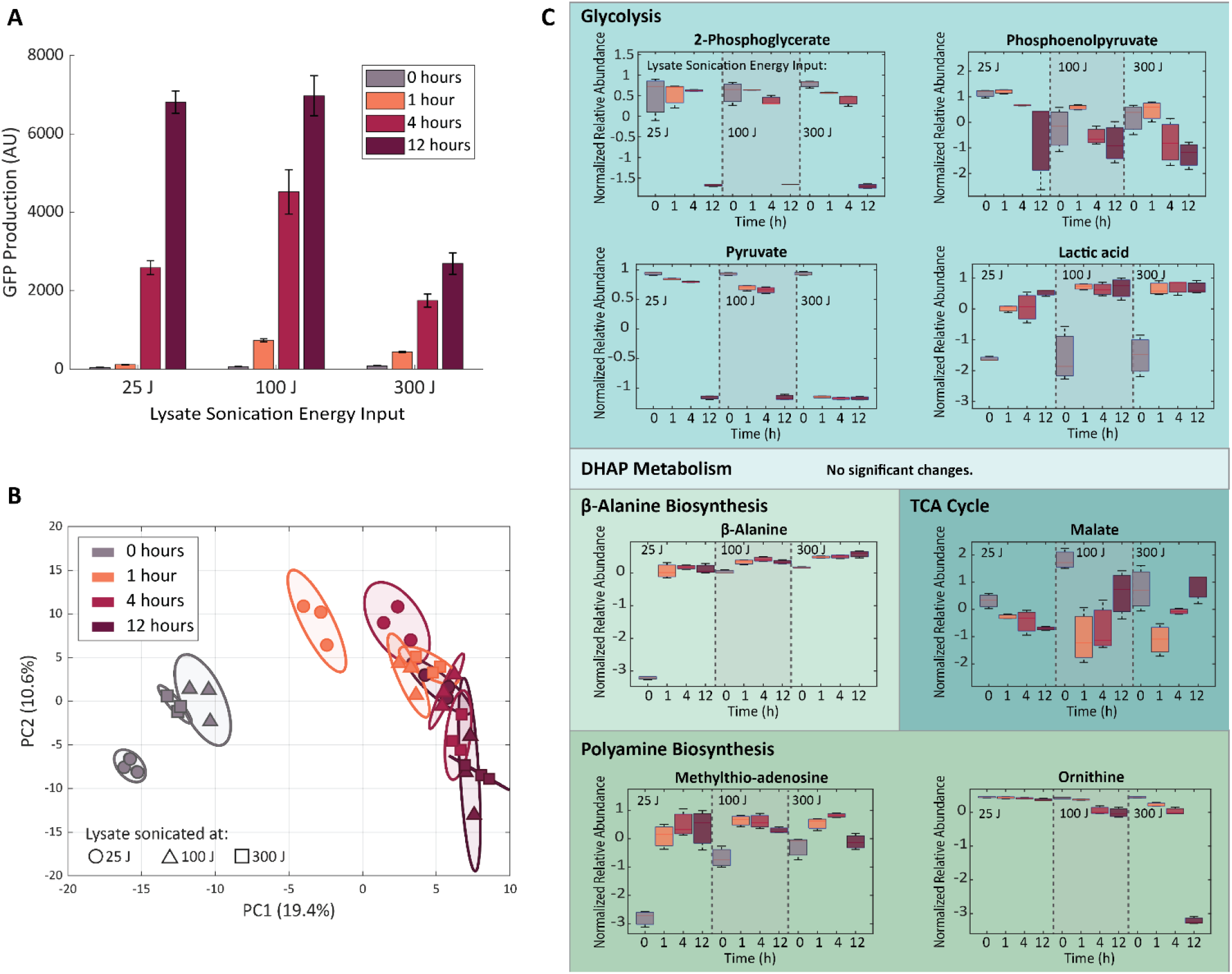
GFP production and metabolic changes in CFE reactions using lysates sonicated with different energy inputs. (A) Reducing lysate sonication energy input from 300 J to 25 or 100 J significantly improves protein production expression in CFE reactions. Error bars represent standard deviation of triplicate reactions. (B) Principal component analysis of metabolite profiles from reactions using differently sonicated lysates. The different lysate reactions separate in PC1 at 0, 1, and 4 h timepoints. Colored ellipses represented 95% confidence intervals for each group, and the plotted samples are replicate reactions. (C) Metabolites involved in glycolysis, the TCA cycle, β-alanine biosynthesis, and polyamine biosynthesis pathways are prominently affected by sonication energy input. Box and whisker plots depict the normalized peak areas, which are transformed using a generalized logarithm (base 2) and autoscaled. Red lines are the medians, boxes span the second and third quartiles of values. Error bars represent standard deviation of triplicate reactions.

Univariate (ANOVA) analysis again provided additional context for the multivariate results. 100 and 300 J reactions behaved like our complete CFE reactions and had almost identical metabolic behavior, with few exceptions **(Figure 5C)**. The 25 J reactions appeared to have slower glycolytic activity, as evidenced by the smaller changes between 0 h and 4 h compared to the 100 J and 300 J reactions. Interestingly, the 25 J reactions had lower initial abundances of malate, β-alanine, and methylthio-adenosine, although the trends of all but malate remained the same across sonication energies. Glycerol-3-phosphate, fumarate, succinate, and putrescine were present in the data set and remained constant for all reactions **(Figure S7A-D)**.

ANOVA2 results highlight the importance of time and interaction effects over sonication energy input alone, indicating that (similar to lysate pre-incubation) the initial metabolic differences between lysates with different sonication energies are ultimately blunted over time **(Figure S8A).** While the quantitative metabolic dynamics across these lysates differ, they are often variations on similar trends but to different degrees. This may be a result of the change in total protein content (rather than specific activity) caused by altering sonication energy input. Even though the lysates for the sonication energy input experiment were derived from a different strain and prepared by a different operator than in the previous figures, their metabolic dynamics were similar to the previous results; this further highlights the resiliency of CFE reaction metabolism to external perturbations, which is a particularly salient feature given the well-known operator dependence of CFE results.

We note that because the stability and function of proteins are known to be affected by sonication energy input, proteins involved in transcription and translation could have been affected in our experiment, though they were not directly measured. However, even if these effects are present, they seem to be dominated by other effects of changing sonication energies (e.g., degree of cell lysis and permeabilization). The metabolic behaviors observed here, along with the unique temporal expression dynamics at different sonication energies, indicate the likely complex interdependencies between proteins and metabolites, and between expression and metabolism, in CFE systems.

### Targeted enzyme supplementation has minor effects on protein yield and metabolic profiles

With this additional support for the idea that endogenous metabolism is connected to CFE productivity and potentially robust to changes in the initial lysate’s metabolic state, we sought to assess the impact on productivity when some metabolic fluxes are perturbed or rerouted, which could perhaps cause larger-scale metabolic perturbations than single metabolite supplementation. We selected five cytosolic enzymes to supplement in CFE reactions based on our findings to this point and their known importance in central carbon and amino acid metabolism: three glycolytic enzymes (phosphoglycerate kinase (*pgk*), pyruvate kinase (*pykF*), and lactate dehydrogenase (*ldhA*)) and two TCA cycle enzymes (citrate synthase (*gltA*) and isocitrate dehydrogenase (*icd*)) **(Figure 1)**.

We initially sought to supplement the enzymes via expression from a plasmid; however, we found that including additional plasmid DNA to produce the enzyme at the same time as the GFP reporter in a CFE reaction confounded results **(Figure S9)**. Extra DNA, regardless of sequence and gene products, increased GFP production in these experiments. To avoid this confounding effect, we instead supplemented the reaction with purified enzyme. Before performing metabolomics analysis, we identified the optimal concentration ranges for expression enhancement for each enzyme in small-volume reactions **(Figure S10).** Only two of the five enzymes (GltA and LdhA) caused improved endpoint GFP production in the CFE reactions when supplemented at their optimal concentrations.

We selected GltA, Pgk, and LdhA for metabolomics analysis of supplemented reactions based on their metabolic pathway diversity (they come from the TCA cycle, glycolysis, and fermentation, respectively) and the preliminary evidence of improved GFP production for GltA and LdhA. Each larger-volume reaction was supplemented with the optimized concentration of the selected enzyme (1 μM GltA, 100 nM LdhA,10 nM Pgk); control reactions were supplemented with enzyme buffer with no enzyme. Samples were collected for fluorescence and metabolomics analysis at 0, 1, and 4 hours after reaction assembly. After data processing, these metabolomics measurements yielded 331 known and unannotated analytes used in further analyses.

Only GltA supplementation yielded significantly improved GFP expression in the large-volume reactions **(Figure 6A)**, though supplementation with the other enzymes trended in the same direction. (Differences in protein expression between small- and large-volume reactions is consistent with literature reports^29^.) Multivariate analysis via PCA yielded no clear separation of samples at each timepoint (though the GltA samples showed a small amount of separation at 0 hours), indicating that supplementation of these enzymes only minimally impacts the metabolic profile despite their known importance in carbon metabolism **(Figure 6B)**.

**Figure 6.**
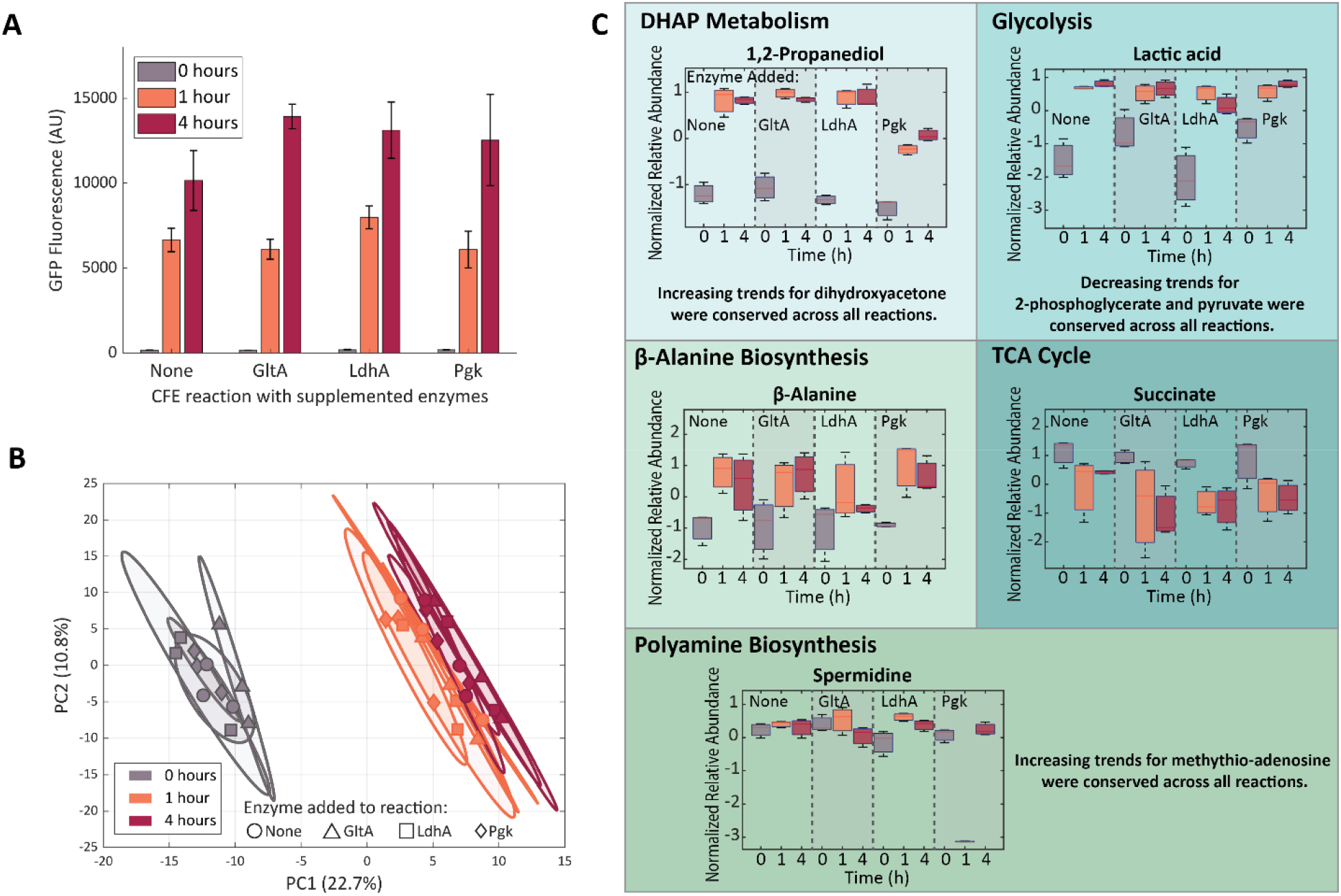
GFP production and metabolic changes in CFE reactions supplemented with the enzymes GltA, LdhA, or Pgk. (A) Though all three supplemented reactions trended towards increased GFP expression, only GltA supplementation yielded a statistically significant increase at 4 h. Error bars represent standard deviation of triplicate reactions. (B) Principal component analysis of metabolite profiles collected at different reaction timepoints. The supplementation conditions do not clearly separate at any timepoint. Colored ellipses represent 95% confidence intervals for each group, and the plotted samples are replicate reactions. (C) Only a few metabolites involved in glycolysis, DHAP metabolism, the TCA cycle, β-alanine biosynthesis, and polyamine biosynthesis pathways were affected by enzyme supplementation. Box and whisker plots depict the normalized peak areas, which are transformed using a generalized logarithm (base 2) and autoscaled. Red lines are the medians, boxes span the second and third quartiles of values. Error bars represent standard deviation of triplicate reactions.

On a univariate level, while most metabolite levels were not affected by enzyme supplementation, there were five metabolites within DHAP metabolism, glycolysis, TCA cycle, β-alanine production and polyamine biosynthesis with statistically significant (though often subtle) changes for different supplemented enzymes (**Figure 6C**). The DHAP breakdown product 1,2-propanediol increased for all reactions, but to a lesser extent for reactions with Pgk, potentially indicating some rerouting of metabolic flux from DHAP metabolism into glycolysis. Reactions with LdhA had lower levels of lactic acid at 4 hours than the control reaction. Although one may have expected larger changes to lactic acid production for reactions supplemented with LdhA, some of the lactic acid pool may come from conversion of methylglyoxal in DHAP metabolism. Succinate levels also change with enzyme supplementation (though with similar profiles across all three enzymes), and supplementation of LdhA and Pgk both affected biosynthesis of the polyamine spermidine. For GltA supplementation (again, the only one to yield significant increases in GFP expression), only one metabolite (succinate) was significantly changed compared to the control. Interestingly, β-alanine levels were largely unaffected, even though one might have anticipated changes based on GltA’s consumption of OAA. β-alanine profiles may have remained unchanged due to the excess of amino acids (specifically L-aspartate) supplied in the reaction mixture.

Taken together, the impacts of enzyme supplementation on the metabolic state of the CFE reactions is comparatively small (though again, one should note the caveat that GC-MS is not well-suited to measuring all classes of metabolites, so there may metabolites we did not measure with more clear differentiation between conditions). While analysis with ANOVA2 suggests that there are more statistically significant enzyme supplementation effects than what is evident from visual inspection or one-way analyses, time still has the broadest effect on metabolite profiles **(Figure S11A)** and the ANOVA2-significant enzyme effects are still quite subtle. Although one may have expected more substantial changes to metabolic activity due to enzyme supplementation, our results suggest a certain degree of resilience of CFE metabolism, perhaps a result of the allosteric regulation that allows living *E. coli* cells to maintain metabolite homeostasis^30^. This overall metabolic resilience only further highlights the importance of using metabolism as a guide or target for optimization of CFE systems, as it is a prominent force with a substantial impact on total protein expression.

## CONCLUSION

In this work, we identified temporal changes in the small molecules within a CFE reaction via metabolomics using GC×GC-MS analysis, linking them to key areas of metabolism. We dissected the contributions of the lysate and reaction mixture to the metabolic changes in a complete CFE reaction, confirming that the endogenous metabolic activity of the lysate plays a significant role. We also demonstrated that lysate pre-incubation and changes to lysate sonication energy input alter protein yield putatively via metabolic changes but have comparatively minor impacts on the overall metabolic profile of the reaction, suggesting that subtle and small changes in metabolite levels may play a significant role in determining reaction productivity. Furthermore, selectively targeting some of the most affected areas of metabolism via enzyme supplementation was able to improve expression, though only in a few cases and it had only minor impact on the metabolic profile of the reaction. Overall, CFE reactions maintain a robust balance of metabolites despite changes to the initial metabolic state of the lysate and enzymatic capacity of the lysate. Our results highlight the complex, intertwined relationship of metabolism and protein expression and their potential as a vehicle for both understanding and optimization of CFE systems.

## Methods

### Plasmid Preparation

The plasmids pJL1s70 and E01 were used in this study. The sequences of both plasmids are available in the c. Plasmids were transformed into *E. coli* DH10B cells and isolated with E.Z.N.A FastFilter Plasmid Maxiprep kit (Omega Biotek) according to the manufacturer’s instructions.

### Lysate Preparation

Cellular lysate for all experiments was prepared based on previously described protocols^28^. Briefly, BL21 cells were used for all experiments, except for the experiment evaluating the effects of sonication energy input, where BL21 DE3 Star Δ*lacZ* cells were used. All cells were grown in either 2× YTP media (16 g L^−1^ tryptone, 10 g L^−1^ yeast extract, 5 g L^−1^ sodium chloride, 7 g L^−1^ potassium phosphate dibasic, and 3 g L^−1^ potassium phosphate monobasic and was pH-corrected to 7.2 with Tris base). All media was filter-sterilized prior to use. Cells were grown at 37 °C and 180 rpm to an OD of 2.0, which corresponds with the mid exponential growth phase. Cells were then centrifuged at 2,700 rcf and washed three times with S30A buffer (14 mM magnesium acetate, 60 mM potassium acetate, 10 mM Tris-acetate (pH 8.2), and 2 mM dithiothreitol). After the final centrifugation, the wet cell mass was determined, and cells were resuspended in 1 mL of S30A buffer per 1 g of wet cell mass. The cellular resuspension was divided into 1 mL aliquots. Cells were lysed using a Q125 Sonicator (Qsonica, Newton, CT) at a frequency of 20 kHz and at 50% of amplitude. Cells were sonicated on ice with three cycles of 10 s on, 10 s off, delivering approximately 300 J unless otherwise specified in text. An additional 4 mM of dithiothreitol was added to each tube, and the sonicated mixture was then centrifuged at 12000 rcf and 4 °C for 10 min. For lysates prepared with BL21 cells, the supernatant was removed and divided into 1 mL aliquots for run-off reaction at 37 °C and 180 rpm for 80 min. After this runoff reaction, the cellular lysate was centrifuged at 12 000 rcf and 4 °C for 10 min. The supernatant was removed and loaded into a 10 kDa MWCO dialysis cassette (Thermo Scientific). Lysate was dialyzed in 1L of S30B buffer (14 mM magnesium glutamate, 60 mM potassium glutamate, 1 mM dithiothreitol, pH-corrected to 8.2 with Tris) at 4 °C for 3 hours. Dialyzed lysate was removed and centrifuged at 12 000 rcf and 4 °C for 10 min. The supernatant was removed, aliquoted, and stored at −80 °C for future use. For lysates prepared with BL21 DE3 Star Δ*lacZ* cells, the supernatant was removed and divided into 0.5 mL aliquots for run-off reaction at 37 °C and 220 rpm for 80 min. All downstream processing steps were the same as those for lysates prepared with BL21 cells, with the exception of 500 mL of S30B buffer used during dialysis.

### Protein Purification

Plasmids coding for expression of different his-tagged proteins (sequences provided in the Supporting Information) were transformed into BL21 (DE3) cells and plated on LB plates supplemented with kanamycin to grow overnight. One colony was selected the next day and resuspended in a 50 mL LB culture supplemented with kanamycin for overnight growth. The overnight culture was then diluted 100-fold in 500 mL of fresh 2×YTP media containing kanamycin the next morning and grown until its OD600 reached between 0.4-0.6, at which point 0.4 mM of IPTG was added to induce T7 polymerase expression and thus plasmid-driven protein production. The induced culture was transferred to a shaking water bath for incubation at 30 °C and 180 rpm for 16 hours, when cells were pelleted, weighed, and frozen at −80 °C for storage until cell lysis.

1 g of frozen cell pellet was resuspended in 2 mL of lysis buffer (50 mM Na_2_HPO_4_, 500 mM NaCl, 10 mM imidazole, pH 8). The resuspension was divided into 1 mL aliquots and sonicated until cells appeared visible lysed. Sonicated products were centrifuged at 12 000 rcf and 4 °C for 15 min before purifying on a HisPur Ni-NTA column (Thermo Scientific) according to the manufacturer protocol. Purification was verified by SDS-PAGE. The eluted proteins were loaded into 10 kDa MWCO dialysis cassettes (Thermo Scientific) and dialyzed overnight in the storage buffer (50 mM Tris-HCl pH 7.5, 100 mM NaCl, 1 mM DTT, 1 mM EDTA, 2% DMSO). Following dialysis, proteins were centrifuged at 12 000 rcf at 4 °C for 10 min to remove insoluble fractions. The supernatant was removed, and protein concentration was measured on a Nanodrop 2000 before sub-aliquoting and storage at −20 °C.

### CFE Reactions

Cell-free reactions for all experiments were run as previously described^28^. Each cell-free reaction contained 0.85 mM each of GTP, UTP, and CTP, in addition to 1.2 mM ATP, 34 μg/mL of folinic acid, 170 μg/mL E. coli tRNA mixture, 130 mM potassium glutamate, 10 mM ammonium glutamate, 12 mM magnesium glutamate, 2 mM each of the 20 standard amino acids, 0.33 mM nicotine adenine dinucleotide (NAD), 0.27 mM coenzyme-A (CoA), 1.5 mM spermidine, 1 mM putrescine, 4 mM sodium oxalate, 33 mM phosphoenolpyruvate (PEP), 27% cell lysate, and 12 nM of the specified plasmid. (9 nM of pJL1s70 was used for the experiment evaluating the effects of sonication energy input.)

For metabolomics analysis, 210 μL reactions were prepared in 1.5 mL microcentrifuge tubes in technical triplicates. Samples were incubated at 37 °C for the specified time. A total of 10 μL of the reaction was then removed and stored at −80 °C for subsequent fluorescence analysis on a BioTek Synergy H4 microplate reader (485/510 nm excitation/emission wavelength, gain of 70). and the remaining 200 μL was stored at −80 °C for subsequent metabolomics analysis. In experiments solely assessing GFP production, 10 μL reactions were prepared in technical triplicates in 384-well plates (Greiner Bio-One) and fluorescence values were measured every 5 minutes at 37 °C. A transparent film was used to seal the plates to prevent reagent evaporation.

### Protein Precipitation for Metabolomics Analysis

Before beginning the protein precipitation protocol, a small volume was removed from all samples in an individual experiment to prepare pooled quality control (QC) samples for the mass spectrometry data acquisition. 25 μL was removed from each sample from the in-depth time course analysis of CFE reactions and the comparison of reactions with differently sonicated lysates. 20 μL was removed from each sample for the time course analysis of the incubated lysate and the comparison of enzyme-supplemented reactions. 15.4 and 10 μL were removed from each samples for the time course analysis of the incubated reaction mix and the comparison of reactions with lysates pre-incubated at different temperatures, respectively. These pooled QC samples were prepared with all other samples for protein precipitation.

Proteins were precipitated from all samples stored for metabolomics analysis via the following protocol^31^: first, methanol was added to each sample at a 1:2 sample to methanol ratio and vortexed briefly. The samples were incubated at −20 °C for 20 min and centrifuged at 11 600 rcf for 30 min, and the supernatant was collected. The supernatants of pooled QC samples were then evenly aliquoted into multiple tubes as needed: two tubes for the in-depth time course analysis of CFE reactions and the time course analysis of the incubated lysate; three tubes for the comparison of reactions with lysates pre-incubated at different temperatures, the comparison of enzyme-supplemented reactions, and the comparison of reactions with differently sonicated lysates; and one tube for the time course analysis of the incubated reaction mix. The supernatants were dried at 40 °C in a CentriVap until all water was removed, and stored at −80°.

### GC-MS Analysis

Before derivatization, stored samples were transferred to a CentriVap to be dried at 40 °C for 15 min. Samples were derivatized as previously described^32, 33^. A total of 10 μL of 20 mg/mL *O*-methylhydroxylamine hydrochloride (MP Biomedicals, LLC, Santa Ana, CA, U.S.A.) in pyridine was added to each dried sample and shaken at 1400 rpm for 90 min at 30 °C. A total of 90 μL of N-methyl-N-(trimethylsilyl) trifluoroacetamide (MSTFA) + 1% trimethylchlorosilane (TMCS) (Thermo Scientific, Lafayette, CO, U.S.A.) was then added to the samples and shaken at 1400 rpm for 30 min at 37 °C. Samples were centrifuged at 21 100 rcf for 3 min, and 50 μL of the supernatant was added to an autosampler vial. Samples were spiked with 0.25 μL of a retention time standard solution composed of fatty acid methyl esters (FAMES). At the beginning of the GC-MS run, the QCs were injected once, and this was repeated again after every 4−6 sample injections to allow for downstream correction for batch effects. A derivatization blank was prepared and run with every batch of samples. A LECO Pegasus 4D instrument with an Agilent 7683B autosampler, Agilent 7890A gas chromatograph, and time-of-flight mass spectrometer (TOF-MS) was used to analyze the samples. The first column was an HP-5, 28 m long × 0.320 mm ID × 0.25 μm film thickness (Agilent, Santa Clara, CA, U.S.A.), and the second was an Rtx-200, 1.5 – 1.8 m long × 0.25 mm ID × 0.25 μm film thickness (Restek, Bellefonte, PA, U.S.A.). More detailed gas chromatography, autosampler, and mass spectrometry methods are provided in the Supporting Information.

### Data Analysis

Sample runs were analyzed in ChromaTOF (LECO, St. Joseph, MI, U.S.A.) to determine baseline, peak area, and peak identification as described previously^20, 34^. Briefly, settings included a baseline offset of 0.5, automatic smoothing, first dimension peak width of 36 s, second dimension peak width of 0.10 s, and a match of 700 required to combine peaks with a minimum signal-to-noise (S/N) of 5 for all subpeaks. Peaks were required to have a S/N of 10 and have a minimum similarity score of 800 to NIST, Golm, and in-house spectral libraries before assigning a name. Unique mass was used for area and height calculation. MetPP was used to align the samples^35^. Sample files and a derivatization reagent blank file were uploaded from ChromaTOF. Unknowns were retained during the peak alignment process. The derivatization reagent blank file was used to subtract peaks resulting from the sample preparation reagents from the corresponding sample files. On-the-fly alignment was used with manually selected quality control samples as the peak list for primary alignment. Peak alignment was performed using the default criteria. For the experiment with CFE reactions with differently sonicated lysates, one of the samples from the group with the lysate sonicated at 300 J at 4 hours was not properly injected into the GC-MS. For this group, we used the average of the abundances of the two properly injected samples to allow three samples in the group for compatibility with downstream analysis software.

To remove analytes that were not reproducibly detected, analytes for which more than half of the values were missing in the QC samples or for which the QC samples had a coefficient of variance larger than 0.5 were removed from the data set. Then, missing values were manually corrected using small value correction only if all the values were missing in the biological replicates.

Finally, MetaboAnalyst was used for statistical and two-factor analysis^36^. For both analyses, remaining missing values were k- nearest neighbors (KNN) corrected. Data was then log-transformed using a generalized logarithm (base 2) and autoscaled. *P*-values were adjusted using the Benjamini-Hochberg False Discovery Rate (FDR). Differences were considered significant at FDR-corrected *p*-values < 0.05.

The metabolomics data sets for this study are available via the Metabolights repository, with the data set identifier MTBLS2630^37^.

## Notes

### Competing Interest Statement

The authors have declared no competing interest.

## References

(1) Sun, J. and H.S. Alper, Metabolic engineering of strains: from industrial-scale to lab-scale chemical production. Journal of Industrial Microbiology and Biotechnology, 2015. 42(3): p. 423–436.

(2) McNerney, M.P. and M.P. Styczynski, Precise control of lycopene production to enable a fast-responding, minimal-equipment biosensor. Metab Eng, 2017. 43(Pt A): p. 46–53.

(3) McNerney, M.P., D.M. Watstein, and M.P. Styczynski, Precision metabolic engineering: The design of responsive, selective, and controllable metabolic systems. Metab Eng, 2015. 31: p. 123–31.

(4) Tan, S.Z. and K.L.J. Prather, Dynamic pathway regulation: recent advances and methods of construction. Current Opinion in Chemical Biology, 2017. 41: p. 28–35.

(5) Burg, J.M., C.B. Cooper, Z. Ye, B.R. Reed, E.A. Moreb, and M.D. Lynch, Large-scale bioprocess competitiveness: the potential of dynamic metabolic control in two-stage fermentations. Current Opinion in Chemical Engineering, 2016. 14: p. 121–136.

(6) Tran, K., C. Gurramkonda, M.A. Cooper, M. Pilli, J.E. Taris, N. Selock, T.-C. Han, M. Tolosa, A. Zuber, C. Peñalber-Johnstone, C. Dinkins, N. Pezeshk, Y. Kostov, D.D. Frey, L. Tolosa, D.W. Wood, and G. Rao, Cell-free production of a therapeutic protein: Expression, purification, and characterization of recombinant streptokinase using a CHO lysate. Biotechnology and Bioengineering, 2018. 115(1): p. 92–102.

(7) McNerney, M.P., F. Piorino, C.L. Michel, and M.P. Styczynski, Active Analyte Import Improves the Dynamic Range and Sensitivity of a Vitamin B12 Biosensor. ACS synthetic biology, 2020. 9(2): p. 402–411.

(8) McNerney, M., C. Michel, K. Kishore, J. Standeven, and M. Styczynski, Dynamic and tunable metabolite control for robust minimal-equipment assessment of serum zinc. Nature Communications, 2019. 10: p. 5514.

(9) Bergquist, P.L., S. Siddiqui, and A. Sunna, Cell-Free Biocatalysis for the Production of Platform Chemicals. Frontiers in Energy Research, 2020. 8(193).

(10) Zemella, A., L. Thoring, C. Hoffmeister, and S. Kubick, Cell-Free Protein Synthesis: Pros and Cons of Prokaryotic and Eukaryotic Systems. Chembiochem, 2015. 16(17): p. 2420–2431.

(11) Kim, D.-M. and J.R. Swartz, Prolonging cell-free protein synthesis with a novel ATP regeneration system. Biotechnology and Bioengineering, 1999. 66(3): p. 180–188.

(12) Kim, D.-M. and J.R. Swartz, Regeneration of adenosine triphosphate from glycolytic intermediates for cell-free protein synthesis. Biotechnology and Bioengineering, 2001. 74(4): p. 309–316.

(13) Calhoun, K.A. and J.R. Swartz, Energizing cell-free protein synthesis with glucose metabolism. Biotechnology and Bioengineering, 2005. 90(5): p. 606–613.

(14) Silverman, A.D., N. Kelley-Loughnane, J.B. Lucks, and M.C. Jewett, Deconstructing Cell-Free Extract Preparation for in Vitro Activation of Transcriptional Genetic Circuitry. ACS Synthetic Biology, 2019. 8(2): p. 403–414.

(15) Shin, J. and V. Noireaux, An E. coli Cell-Free Expression Toolbox: Application to Synthetic Gene Circuits and Artificial Cells. ACS Synthetic Biology, 2012. 1(1): p. 29–41.

(16) Garenne, D., C.L. Beisel, and V. Noireaux, Characterization of the all-E. coli transcription-translation system myTXTL by mass spectrometry. Rapid Communications in Mass Spectrometry, 2019. 33(11): p. 1036–1048.

(17) Garcia, D.C., B.P. Mohr, J.T. Dovgan, G.B. Hurst, R.F. Standaert, and M.J. Doktycz, Elucidating the potential of crude cell extracts for producing pyruvate from glucose. Synthetic Biology, 2018. 3(1).

(18) Foshag, D., E. Henrich, E. Hiller, M. Schäfer, C. Kerger, A. Burger-Kentischer, I. Diaz-Moreno, S.M. García-Mauriño, V. Dötsch, S. Rupp, and F. Bernhard, The E. coli S30 lysate proteome: A prototype for cell-free protein production. New Biotechnology, 2018. 40: p. 245–260.

(19) Miguez, A.M., M.P. McNerney, and M.P. Styczynski, Metabolomics Analysis of the Toxic Effects of the Production of Lycopene and Its Precursors. Frontiers in Microbiology, 2018. 9(760).

(20) Dhakshinamoorthy, S., N.-T. Dinh, J. Skolnick, and M.P. Styczynski, Metabolomics identifies the intersection of phosphoethanolamine with menaquinone-triggered apoptosis in an in vitro model of leukemia. Molecular BioSystems, 2015. 11(9): p. 2406–2416.

(21) Miguez, A., M. McNerney, and M. Styczynski, Metabolic Profiling of Escherichia coli – Based Cell-Free Expression Systems for Process Optimization. Industrial & Engineering Chemistry Research, 2019. 58.

(22) Subedi, K.P., I. Kim, J. Kim, B. Min, and C. Park, Role of GldA in dihydroxyacetone and methylglyoxal metabolism of Escherichia coli K12. FEMS Microbiology Letters, 2008. 279(2): p. 180–187.

(23) Ferguson, G.P., S. Tötemeyer, M.J. MacLean, and I.R. Booth, Methylglyoxal production in bacteria: suicide or survival? Archives of microbiology, 1998. 170(4): p. 209–18.

(24) Pontrelli, S., R.C.B. Fricke, S.T. Teoh, W.A. Laviña, S.P. Putri, S. Fitz-Gibbon, M. Chung, M. Pellegrini, E. Fukusaki, and J.C. Liao, Metabolic repair through emergence of new pathways in Escherichia coli. Nature Chemical Biology, 2018. 14(11): p. 1005–1009.

(25) Leonardi, R. and S. Jackowski, Biosynthesis of Pantothenic Acid and Coenzyme A. EcoSal Plus, 2007. 2(2): p. 10.1128/ecosalplus.3.6.3.4.

(26) Guerra, P.R., A. Herrero-Fresno, V. Ladero, B. Redruello, T.P. dos Santos, M.R. Spiegelhauer, L. Jelsbak, and J.E. Olsen, Putrescine biosynthesis and export genes are essential for normal growth of avian pathogenic Escherichia coli. BMC Microbiology, 2018. 18(1): p. 226.

(27) Schwarz, D., F. Junge, F. Durst, N. Frölich, B. Schneider, S. Reckel, S. Sobhanifar, V. Dötsch, and F. Bernhard, Preparative scale expression of membrane proteins in Escherichia coli-based continuous exchange cell-free systems. Nature Protocols, 2007. 2(11): p. 2945–2957.

(28) Kwon, Y.-C. and M.C. Jewett, High-throughput preparation methods of crude extract for robust cell-free protein synthesis. Scientific Reports, 2015. 5(1): p. 8663.

(29) Voloshin, A.M. and J.R. Swartz, Efficient and scalable method for scaling up cell free protein synthesis in batch mode. Biotechnol Bioeng, 2005. 91(4): p. 516–21.

(30) Yasid, N.A., M.D. Rolfe, J. Green, and M.P. Williamson, Homeostasis of metabolites in Escherichia coli on transition from anaerobic to aerobic conditions and the transient secretion of pyruvate. R Soc Open Sci, 2016. 3(8): p. 160187–160187.

(31) Gowda, G.A.N. and D. Raftery, Quantitating Metabolites in Protein Precipitated Serum Using NMR Spectroscopy. Analytical Chemistry, 2014. 86(11): p. 5433–5440.

(32) Kind, T., G. Wohlgemuth, D.Y. Lee, Y. Lu, M. Palazoglu, S. Shahbaz, and O. Fiehn, FiehnLib: Mass Spectral and Retention Index Libraries for Metabolomics Based on Quadrupole and Time-of-Flight Gas Chromatography/Mass Spectrometry. Analytical Chemistry, 2009. 81(24): p. 10038–10048.

(33) Vermeersch, K.A., L. Wang, J.F. McDonald, and M.P. Styczynski, Distinct metabolic responses of an ovarian cancer stem cell line. BMC Systems Biology, 2014. 8(1): p. 134.

(34) Vermeersch, K.A., L. Wang, R. Mezencev, J.F. McDonald, and M.P. Styczynski, OVCAR-3 Spheroid-Derived Cells Display Distinct Metabolic Profiles. PLOS ONE, 2015. 10(2): p. e0118262.

(35) Wei, X., X. Shi, I. Koo, S. Kim, R.H. Schmidt, G.E. Arteel, W.H. Watson, C. McClain, and X. Zhang, MetPP: a computational platform for comprehensive two-dimensional gas chromatography time-of-flight mass spectrometry-based metabolomics. Bioinformatics, 2013. 29(14): p. 1786–1792.

(36) Chong, J., D.S. Wishart, and J. Xia, Using MetaboAnalyst 4.0 for Comprehensive and Integrative Metabolomics Data Analysis. Current protocols in bioinformatics, 2019. 68(1): p. e86.

(37) Haug, K., K. Cochrane, V.C. Nainala, M. Williams, J. Chang, K.V. Jayaseelan, and C. O’Donovan, MetaboLights: a resource evolving in response to the needs of its scientific community. Nucleic Acids Research, 2019. 48(D1): p. D440–D444.

