## Supplementary figures and information for "Metabolic Dynamics in *Escherichia coli*-based Cell-Free Systems"

### GCxGC-MS Methods

#### Auto-sampler Method

An Agilent 7683 auto sampler was used. Prior to sample injection, three pre-washes were performed with pyridine. Samples were pumped 4 times for thorough mixing and injected using a syringe size of 10 µL with 1 µL of injection volume. Three post-washes of the needle were performed after injection using pyridine.

#### GCxGC Method

An Agilent 7890 gas chromatograph adapted to GCxGC analysis was used. Helium was used as the carrier gas with a corrected constant flow rate of 1.00 mL/min. The inlet septum purge flow was maintained at 3 mL/min. The inlet was used in splitless mode with a purge flow of 100 mL/min delayed to start 30 seconds after injection, giving a total flow of 101 mL/min. Runs were performed in a gas saver mode with a flow of 20 mL/min set to start a minute after injection. The front inlet temperature was set at 250°C for the entire run.

The primary oven temperature was held at 70°C for 1 min and the temperature was ramped at 10°C/min until 315°C and held for 2 minutes. The secondary oven temperature and the modulator temperature offsets were 5°C and 15°C above the main oven, respectively. A one-minute equilibration time was set for the ovens. The modulation program is listed in Supplementary Table 1. The transfer line temperature was maintained at 320°C for the entire run.

Table S1. Modulation Timing

| # | Start (s) | End (s) | Modulation period (s) | Hot pulse time (s) | Cool time between stages (s) |
| --- | --- | --- | --- | --- | --- |
| 1 | Start | 392 | 6.00 | 1.00 | 2.00 |
| 2 | 392 | End of Run | 6.00 | 1.50 | 1.50 |

**MS Method**

A Leco Pegasus 4D time of flight mass spectrometer (TOF-MS) with electron impact ionization was used for mass analysis. Filaments were turned off for the initial 230 seconds to delay mass acquisition until after the solvent peak. The mass scanning range was from 50 to 500 u with an acquisition rate of 200 spectra per second. The detector voltage was set at 100 V above the optimized voltage with an electron energy of -70 V. Manual mass defect mode was used with the mass defect 0 mu/ 100 u. The ion source temperature was required to reach 220°C before starting mass acquisition.

### Plasmids used in this study

**pJL1s70:**

agatcaaaggatcttcttgagatcctttttttctgcgcgtaatctgctgcttgcaaacaaaaaaaccaccgctaccagcggtggtttgtttgccggatcaagagctaccaactctttttccgaaggtaactggcttcagcagagcgcagataccaaatactgttcttctagtgtagccgtagttaggccaccacttcaagaactctgtagcaccgcctacatacctcgctctgctaatcctgttaccagtggctgctgccagtggcgataagtcgtgtcttaccgggttggactcaagacgatagttaccggataaggcgcagcggtcgggctgaacggggggttcgtgcacacagcccagcttggagcgaacgacctacaccgaactgagatacctacagcgtgagctatgagaaagcgccacgcttcccgaagggagaaaggcggacaggtatccggtaagcggcagggtcggaacaggagagcgcacgagggagcttccagggggaaacgcctggtatctttatagtcctgtcgggtttcgccacctctgacttgagcgtcgatttttgtgatgctcgtcaggggggcggagcctatggaaaaacgccagcaacgcgatcccgcgaaatttgacggctagctcagtcctaggtacagtgctagcccacaacggtttccctctagaaataattttgtttaactttaagaaggagatatacatATGAGCAAAGGTGAAGAACTGTTTACCGGCGTTGTGCCGATTCTGGTGGAACTGGATGGCGATGTGAACGGTCACAAATTCAGCGTGCGTGGTGAAGGTGAAGGCGATGCCACGATTGGCAAACTGACGCTGAAATTTATCTGCACCACCGGCAAACTGCCGGTGCCGTGGCCGACGCTGGTGACCACCCTGACCTATGGCGTTCAGTGTTTTAGTCGCTATCCGGATCACATGAAACGTCACGATTTCTTTAAATCTGCAATGCCGGAAGGCTATGTGCAGGAACGTACGATTAGCTTTAAAGATGATGGCAAATATAAAACGCGCGCCGTTGTGAAATTTGAAGGCGATACCCTGGTGAACCGCATTGAACTGAAAGGCACGGATTTTAAAGAAGATGGCAATATCCTGGGCCATAAACTGGAATACAACTTTAATAGCCATAATGTTTATATTACGGCGGATAAACAGAAAAATGGCATCAAAGCGAATTTTACCGTTCGCCATAACGTTGAAGATGGCAGTGTGCAGCTGGCAGATCATTATCAGCAGAATACCCCGATTGGTGATGGTCCGGTGCTGCTGCCGGATAATCATTATCTGAGCACGCAGACCGTTCTGTCTAAAGATCCGAACGAAAAAGGCACGCGGGACCACATGGTTCTGCACGAATATGTGAATGCGGCAGGTATTACGTGGAGCCATCCGCAGTTCGAAAAATAAgtcgaccggctgctaacaaagcccgaaaggaagctgagttggctgctgccaccgctgagcaataactagcataaccccttggggcctctaaacgggtcttgaggggttttttgctgaaagccaattctgattagaaaaactcatcgagcatcaaatgaaactgcaatttattcatatcaggattatcaataccatatttttgaaaaagccgtttctgtaatgaaggagaaaactcaccgaggcagttccataggatggcaagatcctggtatcggtctgcgattccgactcgtccaacatcaatacaacctattaatttcccctcgtcaaaaataaggttatcaagtgagaaatcaccatgagtgacgactgaatccggtgagaatggcaaaagcttatgcatttctttccagacttgttcaacaggccagccattacgctcgtcatcaaaatcactcgcatcaaccaaaccgttattcattcgtgattgcgcctgagcgagacgaaatacgcgatcgctgttaaaaggacaattacaaacaggaatcgaatgcaaccggcgcaggaacactgccagcgcatcaacaatattttcacctgaatcaggatattcttctaatacctggaatgctgttttcccggggatcgcagtggtgagtaaccatgcatcatcaggagtacggataaaatgcttgatggtcggaagaggcataaattccgtcagccagtttagtctgaccatctcatctgtaacatcattggcaacgctacctttgccatgtttcagaaacaactctggcgcatcgggcttcccatacaatcgatagattgtcgcacctgattgcccgacattatcgcgagcccatttatacccatataaatcagcatccatgttggaatttaatcgcggcttcgagcaagacgtttcccgttgaatatggctcataacaccccttgtattactgtttatgtaagcagacagttttattgttcatgatgatatatttttatcttgtgcaatgtaacatcagagattttgagacacaacgtg

ColE1 origin

E. coli σ^70^ promoter J23100

5’ UTR

sfGFP

kanamycin resistance marker

**E01:**

agatcaaaggatcttcttgagatcctttttttctgcgcgtaatctgctgcttgcaaacaaaaaaaccaccgctaccagcggtggtttgtttgccggatcaagagctaccaactctttttccgaaggtaactggcttcagcagagcgcagataccaaatactgttcttctagtgtagccgtagttaggccaccacttcaagaactctgtagcaccgcctacatacctcgctctgctaatcctgttaccagtggctgctgccagtggcgataagtcgtgtcttaccgggttggactcaagacgatagttaccggataaggcgcagcggtcgggctgaacggggggttcgtgcacacagcccagcttggagcgaacgacctacaccgaactgagatacctacagcgtgagctatgagaaagcgccacgcttcccgaagggagaaaggcggacaggtatccggtaagcggcagggtcggaacaggagagcgcacgagggagcttccagggggaaacgcctggtatctttatagtcctgtcgggtttcgccacctctgacttgagcgtcgatttttgtgatgctcgtcaggggggcggagcctatggaaaaacgccagcaacgcgatcccgcgaaattaatacgactcactatagggagagtcgaccggctgctaacaaagcccgaaaggaagctgagttggctgctgccaccgctgagcaataactagcataaccccttggggcctctaaacgggtcttgaggggttttttgctgaaagccaattctgattagaaaaactcatcgagcatcaaatgaaactgcaatttattcatatcaggattatcaataccatatttttgaaaaagccgtttctgtaatgaaggagaaaactcaccgaggcagttccataggatggcaagatcctggtatcggtctgcgattccgactcgtccaacatcaatacaacctattaatttcccctcgtcaaaaataaggttatcaagtgagaaatcaccatgagtgacgactgaatccggtgagaatggcaaaagcttatgcatttctttccagacttgttcaacaggccagccattacgctcgtcatcaaaatcactcgcatcaaccaaaccgttattcattcgtgattgcgcctgagcgagacgaaatacgcgatcgctgttaaaaggacaattacaaacaggaatcgaatgcaaccggcgcaggaacactgccagcgcatcaacaatattttcacctgaatcaggatattcttctaatacctggaatgctgttttcccggggatcgcagtggtgagtaaccatgcatcatcaggagtacggataaaatgcttgatggtcggaagaggcataaattccgtcagccagtttagtctgaccatctcatctgtaacatcattggcaacgctacctttgccatgtttcagaaacaactctggcgcatcgggcttcccatacaatcgatagattgtcgcacctgattgcccgacattatcgcgagcccatttatacccatataaatcagcatccatgttggaatttaatcgcggcttcgagcaagacgtttcccgttgaatatggctcataacaccccttgtattactgtttatgtaagcagacagttttattgttcatgatgatatatttttatcttgtgcaatgtaacatcagagattttgagacacaacgtg

ColE1 origin

T7 promoter

kanamycin resistance marker

Proteins sequences used in this study

**P_T7_-his6-gltA**

agatcaaaggatcttcttgagatcctttttttctgcgcgtaatctgctgcttgcaaacaaaaaaaccaccgctaccagcggtggtttgtttgccggatcaagagctaccaactctttttccgaaggtaactggcttcagcagagcgcagataccaaatactgttcttctagtgtagccgtagttaggccaccacttcaagaactctgtagcaccgcctacatacctcgctctgctaatcctgttaccagtggctgctgccagtggcgataagtcgtgtcttaccgggttggactcaagacgatagttaccggataaggcgcagcggtcgggctgaacggggggttcgtgcacacagcccagcttggagcgaacgacctacaccgaactgagatacctacagcgtgagctatgagaaagcgccacgcttcccgaagggagaaaggcggacaggtatccggtaagcggcagggtcggaacaggagagcgcacgagggagcttccagggggaaacgcctggtatctttatagtcctgtcgggtttcgccacctctgacttgagcgtcgatttttgtgatgctcgtcaggggggcggagcctatggaaaaacgccagcaacgcgatcccgcgaaattaatacgactcactatagggagaccacaacggtttccctctagaaataattttgtttaaNctttaagaaggagatatacatatgCATCACCATCACCATCACGCTGATACAAAAGCAAAACTCACCCTCAACGGGGATACAGCTGTTGAACTGGATGTGCTGAAAGGCACGCTGGGTCAAGATGTTATTGATATCCGTACTCTCGGTTCAAAAGGTGTGTTCACCTTTGACCCAGGCTTCACTTCAACCGCATCCTGCGAATCTAAAATTACTTTTATTGATGGTGATGAAGGTATTTTGCTGCACCGCGGTTTCCCGATCGATCAGCTGGCGACCGATTCTAACTACCTGGAAGTTTGTTACATCCTGCTGAATGGTGAAAAACCGACTCAGGAACAGTATGACGAATTTAAAACTACGGTGACCCGTCATACCATGATCCACGAGCAGATTACCCGTCTGTTCCATGCTTTCCGTCGCGACTCGCATCCAATGGCAGTCATGTGTGGTATTACCGGCGCGCTGGCGGCGTTCTATCACGACTCGCTGGATGTTAACAATCCTCGTCACCGTGAAATTGCCGCGTTCCGCCTGCTGTCGAAAATGCCGACCATGGCCGCGATGTGTTACAAGTATTCCATTGGTCAGCCATTTGTTTACCCGCGCAACGATCTCTCCTACGCCGGTAACTTCCTGAATATGATGTTCTCCACGCCGTGCGAACCGTATGAAGTTAATCCGATTCTGGAACGTGCTATGGACCGTATTCTGATCCTGCACGCTGACCATGAACAGAACGCCTCTACCTCCACCGTGCGTACCGCTGGCTCTTCGGGTGCGAACCCGTTTGCCTGTATCGCAGCAGGTATTGCTTCACTGTGGGGACCTGCGCACGGCGGTGCTAACGAAGCGGCGCTGAAAATGCTGGAAGAAATCAGCTCCGTTAAACACATTCCGGAATTTGTTCGTCGTGCGAAAGACAAAAATGATTCTTTCCGCCTGATGGGCTTCGGTCACCGCGTGTACAAAAATTACGACCCGCGCGCCACCGTAATGCGTGAAACCTGCCATGAAGTGCTGAAAGAGCTGGGCACGAAGGATGACCTGCTGGAAGTGGCTATGGAGCTGGAAAACATCGCGCTGAACGACCCGTACTTTATCGAGAAGAAACTGTACCCGAACGTCGATTTCTACTCTGGTATCATCCTGAAAGCGATGGGTATTCCGTCTTCCATGTTCACCGTCATTTTCGCAATGGCACGTACCGTTGGCTGGATCGCCCACTGGAGCGAAATGCACAGTGACGGTATGAAGATTGCCCGTCCGCGTCAGCTGTATACAGGATATGAAAAACGCGACTTTAAAAGCGATATCAAGCGTtaagtcgaccggctgctaacaaagcccgaaaggaagctgagttggctgctgccaccgctgagcaataactagcataaccccttggggcctctaaacgggtcttgaggggttttttgctgaaagccaattctgattagaaaaactcatcgagcatcaaatgaaactgcaatttattcatatcaggattatcaataccatatttttgaaaaagccgtttctgtaatgaaggagaaaactcaccgaggcagttccataggatggcaagatcctggtatcggtctgcgattccgactcgtccaacatcaatacaacctattaatttcccctcgtcaaaaataaggttatcaagtgagaaatcaccatgagtgacgactgaatccggtgagaatggcaaaagcttatgcatttctttccagacttgttcaacaggccagccattacgctcgtcatcaaaatcactcgcatcaaccaaaccgttattcattcgtgattgcgcctgagcgagacgaaatacgcgatcgctgttaaaaggacaattacaaacaggaatcgaatgcaaccggcgcaggaacactgccagcgcatcaacaatattttcacctgaatcaggatattcttctaatacctggaatgctgttttcccggggatcgcagtggtgagtaaccatgcatcatcaggagtacggataaaatgcttgatggtcggaagaggcataaattccgtcagccagtttagtctgaccatctcatctgtaacatcattggcaacgctacctttgccatgtttcagaaacaactctggcgcatcgggcttcccatacaatcgatagattgtcgcacctgattgcccgacattatcgcgagcccatttatacccatataaatcagcatccatgttggaatttaatcgcggcttcgagcaagacgtttcccgttgaatatggctcataacaccccttgtattactgtttatgtaagcagacagttttattgttcatgatgatatatttttatcttgtgcaatgtaacatcagagattttgagacacaacgtg

ColE1 origin

T7 promoter

5’ UTR

6x His

gltA (Accession ID: EG10402)

kanamycin resistance marker

P_T7_-his6-ldhA

agatcaaaggatcttcttgagatcctttttttctgcgcgtaatctgctgcttgcaaacaaaaaaaccaccgctaccagcggtggtttgtttgccggatcaagagctaccaactctttttccgaaggtaactggcttcagcagagcgcagataccaaatactgttcttctagtgtagccgtagttaggccaccacttcaagaactctgtagcaccgcctacatacctcgctctgctaatcctgttaccagtggctgctgccagtggcgataagtcgtgtcttaccgggttggactcaagacgatagttaccggataaggcgcagcggtcgggctgaacggggggttcgtgcacacagcccagcttggagcgaacgacctacaccgaactgagatacctacagcgtgagctatgagaaagcgccacgcttcccgaagggagaaaggcggacaggtatccggtaagcggcagggtcggaacaggagagcgcacgagggagcttccagggggaaacgcctggtatctttatagtcctgtcgggtttcgccacctctgacttgagcgtcgatttttgtgatgctcgtcaggggggcggagcctatggaaaaacgccagcaacgcgatcccgcgaaattaatacgactcactatagggagaccacaacggtttccctctagaaataattttgtttaaNctttaagaaggagatatacatatgCATCACCATCACCATCACaaactcgccgtttatagcacaaaacagtacgacaagaagtacctgcaacaggtgaacgagtcctttggctttgagctggaattttttgactttctgctgacggaaaaaaccgctaaaactgccaatggctgcgaagcggtatgtattttcgtaaacgatgacggcagccgcccggtgctggaagagctgaaaaagcacggcgttaaatatatcgccctgcgctgtgccggtttcaataacgtcgaccttgacgcggcaaaagaactggggctgaaagtagtccgtgttccagcctatgatccagaggccgttgctgaacacgccatcggtatgatgatgacgctgaaccgccgtattcaccgcgcgtatcagcgtacccgtgatgctaacttctctctggaaggtctgaccggctttactatgtatggcaaaacggcaggcgttatcggtaccggtaaaatcggtgtggcgatgctgcgcattctgaaaggttttggtatgcgtctgctggcgttcgatccgtatccaagtgcagcggcgctggaactcggtgtggagtatgtcgatctgccaaccctgttctctgaatcagacgttatctctctgcactgcccgctgacaccggaaaactatcatctgttgaacgaagccgccttcgaacagatgaaaaatggcgtgatgatcgtcaataccagtcgcggtgcattgattgattctcaggcagcaattgaagcgctgaaaaatcagaaaattggttcgttgggtatggacgtgtatgagaacgaacgcgatctattctttgaagataaatccaacgacgtgatccaggatgacgtattccgtcgcctgtctgcctgccacaacgtgctgtttaccgggcaccaggcattcctgacagcagaagctctgaccagtatttctcagactacgctgcaaaacttaagcaatctggaaaaaggcgaaacctgcccgaacgaactggtttaagtcgaccggctgctaacaaagcccgaaaggaagctgagttggctgctgccaccgctgagcaataactagcataaccccttggggcctctaaacgggtcttgaggggttttttgctgaaagccaattctgattagaaaaactcatcgagcatcaaatgaaactgcaatttattcatatcaggattatcaataccatatttttgaaaaagccgtttctgtaatgaaggagaaaactcaccgaggcagttccataggatggcaagatcctggtatcggtctgcgattccgactcgtccaacatcaatacaacctattaatttcccctcgtcaaaaataaggttatcaagtgagaaatcaccatgagtgacgactgaatccggtgagaatggcaaaagcttatgcatttctttccagacttgttcaacaggccagccattacgctcgtcatcaaaatcactcgcatcaaccaaaccgttattcattcgtgattgcgcctgagcgagacgaaatacgcgatcgctgttaaaaggacaattacaaacaggaatcgaatgcaaccggcgcaggaacactgccagcgcatcaacaatattttcacctgaatcaggatattcttctaatacctggaatgctgttttcccggggatcgcagtggtgagtaaccatgcatcatcaggagtacggataaaatgcttgatggtcggaagaggcataaattccgtcagccagtttagtctgaccatctcatctgtaacatcattggcaacgctacctttgccatgtttcagaaacaactctggcgcatcgggcttcccatacaatcgatagattgtcgcacctgattgcccgacattatcgcgagcccatttatacccatataaatcagcatccatgttggaatttaatcgcggcttcgagcaagacgtttcccgttgaatatggctcataacaccccttgtattactgtttatgtaagcagacagttttattgttcatgatgatatatttttatcttgtgcaatgtaacatcagagattttgagacacaacgtg

ColE1 origin

T7 promoter

5’ UTR

6x His

ldhA (Accession ID: B1380)

kanamycin resistance marker

**P_T7_-his6-pykF**

agatcaaaggatcttcttgagatcctttttttctgcgcgtaatctgctgcttgcaaacaaaaaaaccaccgctaccagcggtggtttgtttgccggatcaagagctaccaactctttttccgaaggtaactggcttcagcagagcgcagataccaaatactgttcttctagtgtagccgtagttaggccaccacttcaagaactctgtagcaccgcctacatacctcgctctgctaatcctgttaccagtggctgctgccagtggcgataagtcgtgtcttaccgggttggactcaagacgatagttaccggataaggcgcagcggtcgggctgaacggggggttcgtgcacacagcccagcttggagcgaacgacctacaccgaactgagatacctacagcgtgagctatgagaaagcgccacgcttcccgaagggagaaaggcggacaggtatccggtaagcggcagggtcggaacaggagagcgcacgagggagcttccagggggaaacgcctggtatctttatagtcctgtcgggtttcgccacctctgacttgagcgtcgatttttgtgatgctcgtcaggggggcggagcctatggaaaaacgccagcaacgcgatcccgcgaaattaatacgactcactatagggagaccacaacggtttccctctagaaataattttgtttaaNctttaagaaggagatatacatatgCATCACCATCACCATCACAAAAAGACCAAAATTGTTTGCACCATCGGACCGAAAACCGAATCTGAAGAGATGTTAGCTAAAATGCTGGACGCTGGCATGAACGTTATGCGTCTGAACTTCTCTCATGGTGACTATGCAGAACACGGTCAGCGCATTCAGAATCTGCGCAACGTGATGAGCAAAACTGGTAAAACCGCCGCTATCCTGCTTGATACCAAAGGTCCGGAAATCCGCACCATGAAACTGGAAGGCGGTAACGACGTTTCTCTGAAAGCTGGTCAGACCTTTACTTTCACCACTGATAAATCTGTTATCGGCAACAGCGAAATGGTTGCGGTAACGTATGAAGGTTTCACTACTGACCTGTCTGTTGGCAACACCGTACTGGTTGACGATGGTCTGATCGGTATGGAAGTTACCGCCATTGAAGGTAACAAAGTTATCTGTAAAGTGCTGAACAACGGTGACCTGGGCGAAAACAAAGGTGTGAACCTGCCTGGCGTTTCCATTGCTCTGCCAGCACTGGCTGAAAAAGACAAACAGGACCTGATCTTTGGTTGCGAACAAGGCGTAGACTTTGTTGCTGCTTCCTTTATTCGTAAGCGTTCTGACGTTATCGAAATCCGTGAGCACCTGAAAGCGCACGGCGGCGAAAACATCCACATCATCTCCAAAATCGAAAACCAGGAAGGCCTCAACAACTTCGACGAAATCCTCGAAGCCTCTGACGGCATCATGGTTGCGCGTGGCGACCTGGGTGTAGAAATCCCGGTAGAAGAAGTTATCTTCGCCCAGAAGATGATGATCGAAAAATGTATCCGTGCACGTAAAGTCGTTATCACTGCGACCCAGATGCTGGATTCCATGATCAAAAACCCACGCCCGACTCGCGCAGAAGCCGGTGACGTTGCAAACGCCATCCTCGACGGTACTGACGCAGTGATGCTGTCTGGTGAATCCGCAAAAGGTAAATACCCGCTGGAAGCGGTTTCTATCATGGCGACCATCTGCGAACGTACCGACCGCGTGATGAACAGCCGTCTCGAGTTCAACAATGACAACCGTAAACTGCGCATTACCGAAGCGGTATGCCGTGGTGCCGTTGAAACTGCTGAAAAACTGGATGCTCCGCTGATCGTGGTTGCTACTCAGGGCGGTAAATCTGCTCGCGCAGTACGTAAATACTTCCCGGATGCCACCATCCTGGCACTGACCACCAACGAAAAAACGGCTCATCAGTTGGTACTGAGCAAAGGCGTTGTGCCGCAGCTTGTTAAAGAGATCACTTCTACTGATGATTTCTACCGTCTGGGTAAAGAACTGGCTCTGCAGAGCGGTCTGGCACACAAAGGTGACGTTGTAGTTATGGTTTCTGGTGCACTGGTACCGAGCGGCACTACTAACACCGCATCTGTTCACGTCCTGtaagtcgaccggctgctaacaaagcccgaaaggaagctgagttggctgctgccaccgctgagcaataactagcataaccccttggggcctctaaacgggtcttgaggggttttttgctgaaagccaattctgattagaaaaactcatcgagcatcaaatgaaactgcaatttattcatatcaggattatcaataccatatttttgaaaaagccgtttctgtaatgaaggagaaaactcaccgaggcagttccataggatggcaagatcctggtatcggtctgcgattccgactcgtccaacatcaatacaacctattaatttcccctcgtcaaaaataaggttatcaagtgagaaatcaccatgagtgacgactgaatccggtgagaatggcaaaagcttatgcatttctttccagacttgttcaacaggccagccattacgctcgtcatcaaaatcactcgcatcaaccaaaccgttattcattcgtgattgcgcctgagcgagacgaaatacgcgatcgctgttaaaaggacaattacaaacaggaatcgaatgcaaccggcgcaggaacactgccagcgcatcaacaatattttcacctgaatcaggatattcttctaatacctggaatgctgttttcccggggatcgcagtggtgagtaaccatgcatcatcaggagtacggataaaatgcttgatggtcggaagaggcataaattccgtcagccagtttagtctgaccatctcatctgtaacatcattggcaacgctacctttgccatgtttcagaaacaactctggcgcatcgggcttcccatacaatcgatagattgtcgcacctgattgcccgacattatcgcgagcccatttatacccatataaatcagcatccatgttggaatttaatcgcggcttcgagcaagacgtttcccgttgaatatggctcataacaccccttgtattactgtttatgtaagcagacagttttattgttcatgatgatatatttttatcttgtgcaatgtaacatcagagattttgagacacaacgtg

ColE1 origin

T7 promoter

5’ UTR

6x His

pykF (Accession ID: EG10804)

kanamycin resistance marker

**P_T7_-his6-icd**

agatcaaaggatcttcttgagatcctttttttctgcgcgtaatctgctgcttgcaaacaaaaaaaccaccgctaccagcggtggtttgtttgccggatcaagagctaccaactctttttccgaaggtaactggcttcagcagagcgcagataccaaatactgttcttctagtgtagccgtagttaggccaccacttcaagaactctgtagcaccgcctacatacctcgctctgctaatcctgttaccagtggctgctgccagtggcgataagtcgtgtcttaccgggttggactcaagacgatagttaccggataaggcgcagcggtcgggctgaacggggggttcgtgcacacagcccagcttggagcgaacgacctacaccgaactgagatacctacagcgtgagctatgagaaagcgccacgcttcccgaagggagaaaggcggacaggtatccggtaagcggcagggtcggaacaggagagcgcacgagggagcttccagggggaaacgcctggtatctttatagtcctgtcgggtttcgccacctctgacttgagcgtcgatttttgtgatgctcgtcaggggggcggagcctatggaaaaacgccagcaacgcgatcccgcgaaattaatacgactcactatagggagaccacaacggtttccctctagaaataattttgtttaaNctttaagaaggagatatacatatgCATCACCATCACCATCACGAAAGTAAAGTAGTTGTTCCGGCACAAGGCAAGAAGATCACCCTGCAAAACGGCAAACTCAACGTTCCTGAAAATCCGATTATCCCTTACATTGAAGGTGATGGAATCGGTGTAGATGTAACCCCAGCCATGCTGAAAGTGGTCGACGCTGCAGTCGAGAAAGCCTATAAAGGCGAGCGTAAAATCTCCTGGATGGAAATTTACACCGGTGAAAAATCCACACAGGTTTATGGTCAGGACGTCTGGCTGCCTGCTGAAACTCTTGATCTGATTCGTGAATATCGCGTTGCCATTAAAGGTCCGCTGACCACTCCGGTTGGTGGCGGTATTCGCTCTCTGAACGTTGCCCTGCGCCAGGAACTGGATCTCTACATCTGCCTGCGTCCGGTACGTTACTATCAGGGCACTCCAAGCCCGGTTAAACACCCTGAACTGACCGATATGGTTATCTTCCGTGAAAACTCGGAAGACATTTATGCGGGTATCGAATGGAAAGCAGACTCTGCCGACGCCGAGAAAGTGATTAAATTCCTGCGTGAAGAGATGGGGGTGAAGAAAATTCGCTTCCCGGAACATTGTGGTATCGGTATTAAGCCGTGTTCGGAAGAAGGCACCAAACGTCTGGTTCGTGCAGCGATCGAATACGCAATTGCTAACGATCGTGACTCTGTGACTCTGGTGCACAAAGGCAACATCATGAAGTTCACCGAAGGAGCGTTTAAAGACTGGGGCTACCAGCTGGCGCGTGAAGAGTTTGGCGGTGAACTGATCGACGGTGGCCCGTGGCTGAAAGTTAAAAACCCGAACACTGGCAAAGAGATCGTCATTAAAGACGTGATTGCTGATGCATTCCTGCAACAGATCCTGCTGCGTCCGGCTGAATATGATGTTATCGCCTGTATGAACCTGAACGGTGACTACATTTCTGACGCCCTGGCAGCGCAGGTTGGCGGTATCGGTATCGCCCCTGGTGCAAACATCGGTGACGAATGCGCCCTGTTTGAAGCCACCCACGGTACTGCGCCGAAATATGCCGGTCAGGACAAAGTAAATCCTGGCTCTATTATTCTCTCCGCTGAGATGATGCTGCGCCACATGGGTTGGACCGAAGCGGCTGACTTAATTGTTAAAGGTATGGAAGGCGCAATCAACGCGAAAACCGTAACCTATGACTTCGAGCGTCTGATGGATGGCGCTAAACTGCTGAAATGTTCAGAGTTTGGTGACGCGATCATCGAAAACATGtaagtcgaccggctgctaacaaagcccgaaaggaagctgagttggctgctgccaccgctgagcaataactagcataaccccttggggcctctaaacgggtcttgaggggttttttgctgaaagccaattctgattagaaaaactcatcgagcatcaaatgaaactgcaatttattcatatcaggattatcaataccatatttttgaaaaagccgtttctgtaatgaaggagaaaactcaccgaggcagttccataggatggcaagatcctggtatcggtctgcgattccgactcgtccaacatcaatacaacctattaatttcccctcgtcaaaaataaggttatcaagtgagaaatcaccatgagtgacgactgaatccggtgagaatggcaaaagcttatgcatttctttccagacttgttcaacaggccagccattacgctcgtcatcaaaatcactcgcatcaaccaaaccgttattcattcgtgattgcgcctgagcgagacgaaatacgcgatcgctgttaaaaggacaattacaaacaggaatcgaatgcaaccggcgcaggaacactgccagcgcatcaacaatattttcacctgaatcaggatattcttctaatacctggaatgctgttttcccggggatcgcagtggtgagtaaccatgcatcatcaggagtacggataaaatgcttgatggtcggaagaggcataaattccgtcagccagtttagtctgaccatctcatctgtaacatcattggcaacgctacctttgccatgtttcagaaacaactctggcgcatcgggcttcccatacaatcgatagattgtcgcacctgattgcccgacattatcgcgagcccatttatacccatataaatcagcatccatgttggaatttaatcgcggcttcgagcaagacgtttcccgttgaatatggctcataacaccccttgtattactgtttatgtaagcagacagttttattgttcatgatgatatatttttatcttgtgcaatgtaacatcagagattttgagacacaacgtg

ColE1 origin

T7 promoter

5’ UTR

6x His

icd (Accession ID: EG10489)

kanamycin resistance marker

**P_T7_-his6-pgk**

agatcaaaggatcttcttgagatcctttttttctgcgcgtaatctgctgcttgcaaacaaaaaaaccaccgctaccagcggtggtttgtttgccggatcaagagctaccaactctttttccgaaggtaactggcttcagcagagcgcagataccaaatactgttcttctagtgtagccgtagttaggccaccacttcaagaactctgtagcaccgcctacatacctcgctctgctaatcctgttaccagtggctgctgccagtggcgataagtcgtgtcttaccgggttggactcaagacgatagttaccggataaggcgcagcggtcgggctgaacggggggttcgtgcacacagcccagcttggagcgaacgacctacaccgaactgagatacctacagcgtgagctatgagaaagcgccacgcttcccgaagggagaaaggcggacaggtatccggtaagcggcagggtcggaacaggagagcgcacgagggagcttccagggggaaacgcctggtatctttatagtcctgtcgggtttcgccacctctgacttgagcgtcgatttttgtgatgctcgtcaggggggcggagcctatggaaaaacgccagcaacgcgatcccgcgaaattaatacgactcactatagggagaccacaacggtttccctctagaaataattttgtttaaNctttaagaaggagatatacatatgCATCACCATCACCATCACTCTGTAATTAAGATGACCGATCTGGATCTTGCTGGGAAACGTGTATTTATCCGTGCGGATCTGAACGTACCAGTAAAAGACGGGAAAGTAACCAGCGACGCGCGTATCCGTGCTTCTCTGCCGACCATCGAACTGGCCCTGAAACAAGGCGCAAAAGTGATGGTAACTTCCCACCTGGGTCGTCCTACCGAAGGCGAGTACAACGAAGAATTCTCTCTGCTGCCGGTTGTTAACTACCTGAAAGACAAACTGTCTAACCCGGTTCGTCTGGTTAAAGATTACCTCGACGGCGTTGACGTTGCTGAAGGTGAACTGGTTGTTCTGGAAAACGTTCGCTTCAACAAAGGCGAGAAGAAAGACGACGAAACCCTGTCCAAAAAATACGCTGCACTGTGTGACGTGTTCGTAATGGACGCATTCGGTACTGCTCACCGCGCGCAGGCTTCTACTCACGGTATCGGTAAATTCGCTGACGTTGCGTGCGCAGGCCCGCTGCTGGCAGCTGAACTGGACGCGCTGGGTAAAGCACTGAAAGAACCTGCTCGCCCGATGGTGGCTATCGTTGGTGGTTCTAAAGTATCTACCAAACTGACCGTTCTGGACTCCCTGTCTAAAATCGCTGACCAGCTGATTGTTGGTGGTGGTATCGCTAACACCTTTATCGCGGCACAAGGCCACGATGTGGGTAAATCCCTGTACGAAGCTGACCTGGTTGACGAAGCTAAACGTCTGCTGACCACCTGCAACATCCCGGTTCCGTCTGATGTTCGCGTAGCAACCGAGTTCTCTGAAACTGCACCGGCTACCCTGAAATCTGTTAACGATGTGAAAGCTGACGAGCAGATCCTGGATATCGGTGATGCTTCCGCTCAGGAACTGGCTGAAATCCTGAAGAATGCGAAAACCATTCTGTGGAACGGTCCGGTTGGCGTGTTCGAATTCCCGAACTTCCGCAAAGGTACTGAAATCGTGGCTAACGCTATCGCAGACAGCGAAGCGTTCTCCATCGCTGGCGGCGGCGACACTCTGGCAGCAATCGACCTGTTCGGCATTGCTGACAAAATCTCCTACATCTCCACTGGCGGCGGCGCATTCCTCGAATTCGTGGAAGGTAAAGTACTGCCTGCAGTAGCGATGCTCGAAGAGCGCGCTAAGAAG_taagtcgaccggctgctaacaaagcccgaaaggaagctgagttggctgctgccaccgctgagcaataactagcataaccccttggggcctctaaacgggtcttgaggggttttttgctgaaagccaattctgattagaaaaactcatcgagcatcaaatgaaactgcaatttattcatatcaggattatcaataccatatttttgaaaaagccgtttctgtaatgaaggagaaaactcaccgaggcagttccataggatggcaagatcctggtatcggtctgcgattccgactcgtccaacatcaatacaacctattaatttcccctcgtcaaaaataaggttatcaagtgagaaatcaccatgagtgacgactgaatccggtgagaatggcaaaagcttatgcatttctttccagacttgttcaacaggccagccattacgctcgtcatcaaaatcactcgcatcaaccaaaccgttattcattcgtgattgcgcctgagcgagacgaaatacgcgatcgctgttaaaaggacaattacaaacaggaatcgaatgcaaccggcgcaggaacactgccagcgcatcaacaatattttcacctgaatcaggatattcttctaatacctggaatgctgttttcccggggatcgcagtggtgagtaaccatgcatcatcaggagtacggataaaatgcttgatggtcggaagaggcataaattccgtcagccagtttagtctgaccatctcatctgtaacatcattggcaacgctacctttgccatgtttcagaaacaactctggcgcatcgggcttcccatacaatcgatagattgtcgcacctgattgcccgacattatcgcgagcccatttatacccatataaatcagcatccatgttggaatttaatcgcggcttcgagcaagacgtttcccgttgaatatggctcataacaccccttgtattactgtttatgtaagcagacagttttattgttcatgatgatatatttttatcttgtgcaatgtaacatcagagattttgagacacaacgtg

ColE1 origin

T7 promoter

5’ UTR

6x His

pgk (Accession ID: EG10703)

kanamycin resistance marker

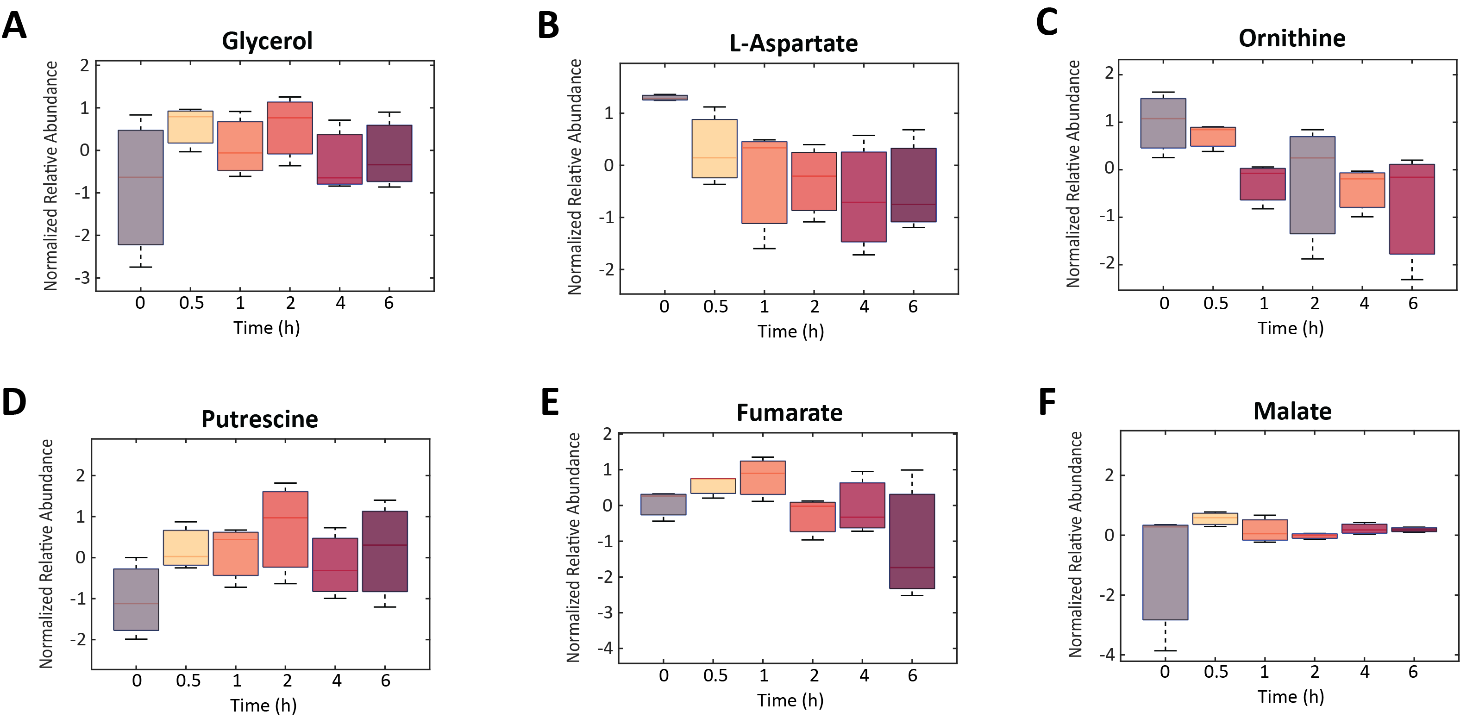
Figure S1. Relative abundances of (A) glycerol, (B) L-aspartate, (C) ornithine, (D) putrescine, (E) fumarate, and (F) malate in CFE reaction samples. Box and whisker plots depict the normalized peak areas, which are transformed using a generalized logarithm (base 2) and autoscaled. Red lines are the medians, boxes represent the second and third quartiles of values. Error bars represent standard deviation of triplicate reactions.

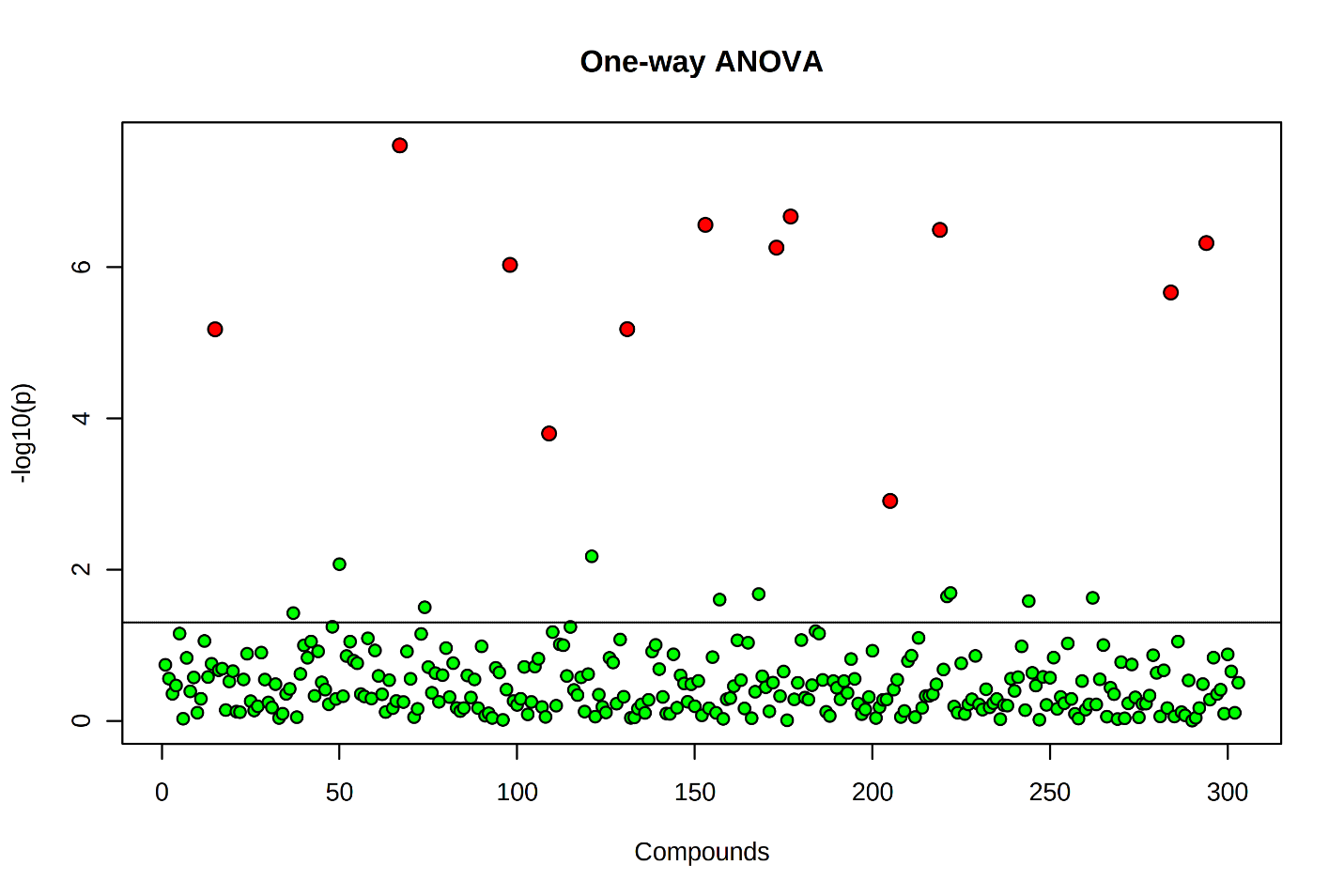

Figure S2. One-way ANOVA of metabolomics data collected from the incubated reaction mixture samples. Incubation of the reaction mixture in water impacted 12 out of 303 analytes detected. Red circles indicate significantly changing analytes, while green circles represent insignificantly changing analytes. The y-axis indicates raw *p*-values for significance tests for each analyte; the *p* = 0.05 line is drawn. Analyte significance was assessed using a false discovery rate (FDR) corrected *p*-value threshold of 0.05.

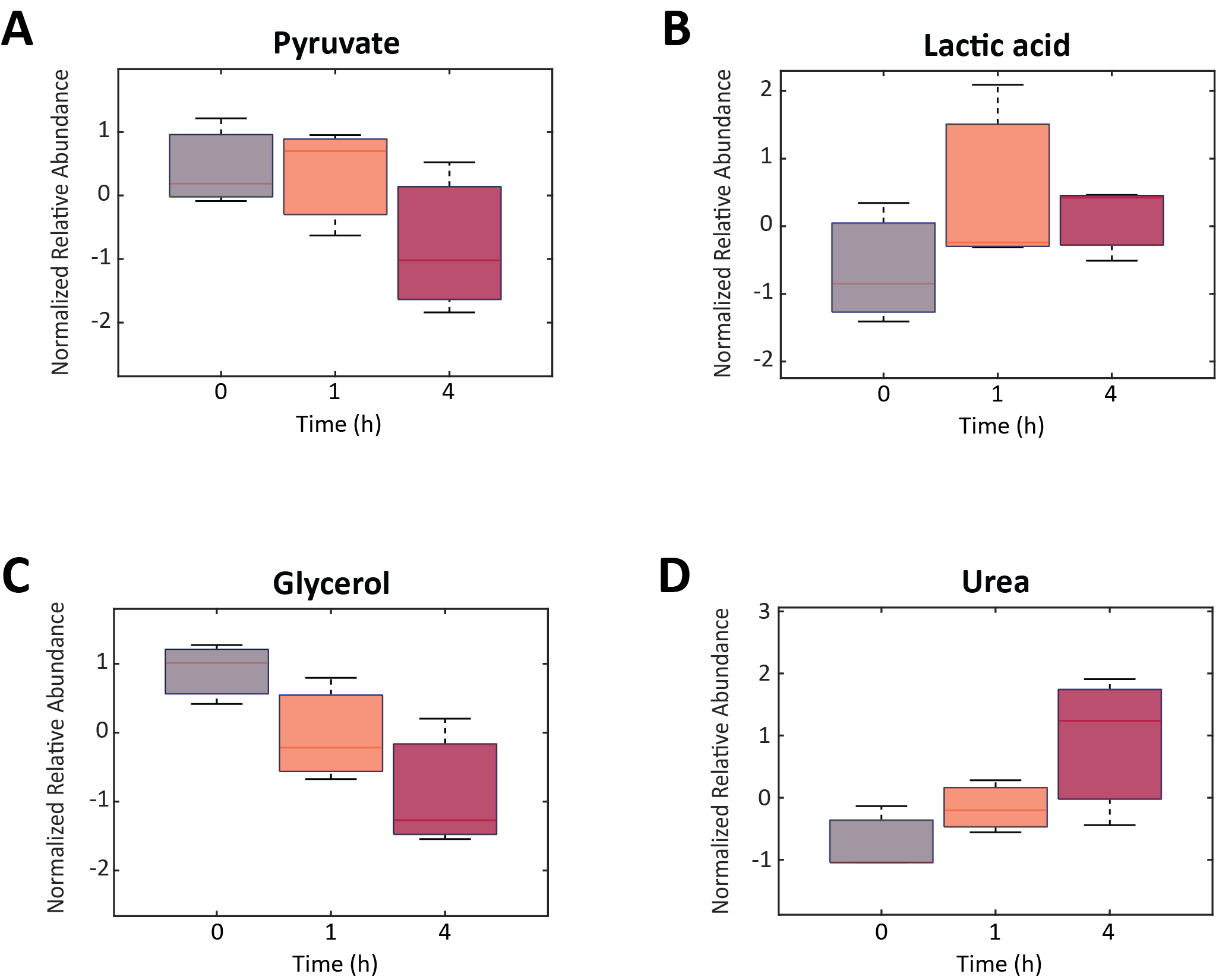

Figure S3. Relative abundances of (A) pyruvate, (B) lactic acid, (C) glycerol, and (D) urea in incubated lysate samples (without reaction mixture or plasmid). Box and whisker plots depict the normalized peak areas, which were transformed using a generalized logarithm (base 2) and autoscaled. Red lines are the medians, boxes are the second and third quartile of values. Error bars represent standard deviation of triplicate reactions.

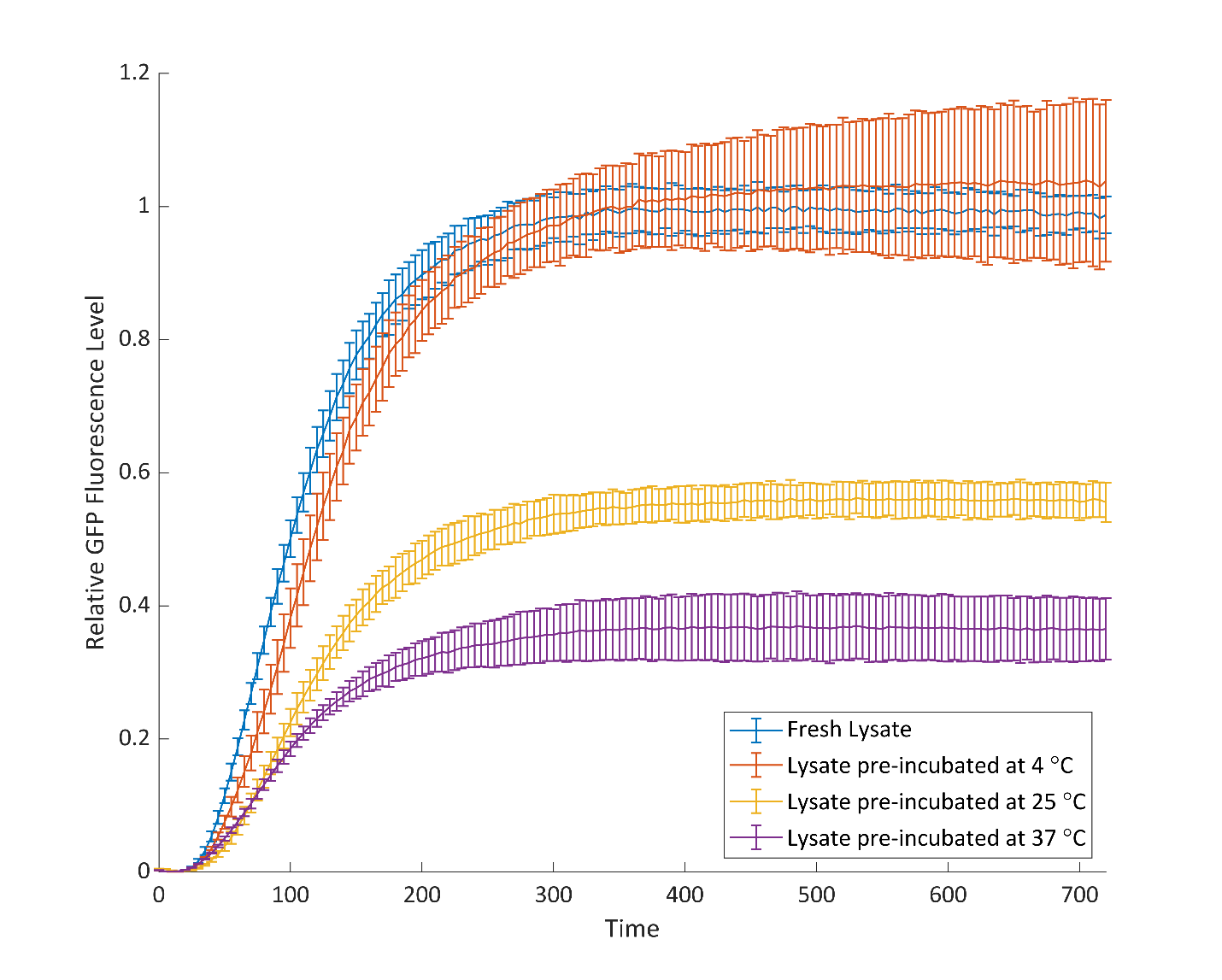
Figure S4. GFP production from CFE reactions using lysates that were fresh or pre-incubated for 6 h at 4, 25, or 37 °C. Fluorescence was measured every 5 minutes over 10 h of incubation with excitation and emission wavelengths of 485 and 510 nm, respectively, with a gain of 70. Each reaction was 10µL in volume with 12nM pJL1s70 plasmid. Error bars represent standard deviation of triplicate reactions.

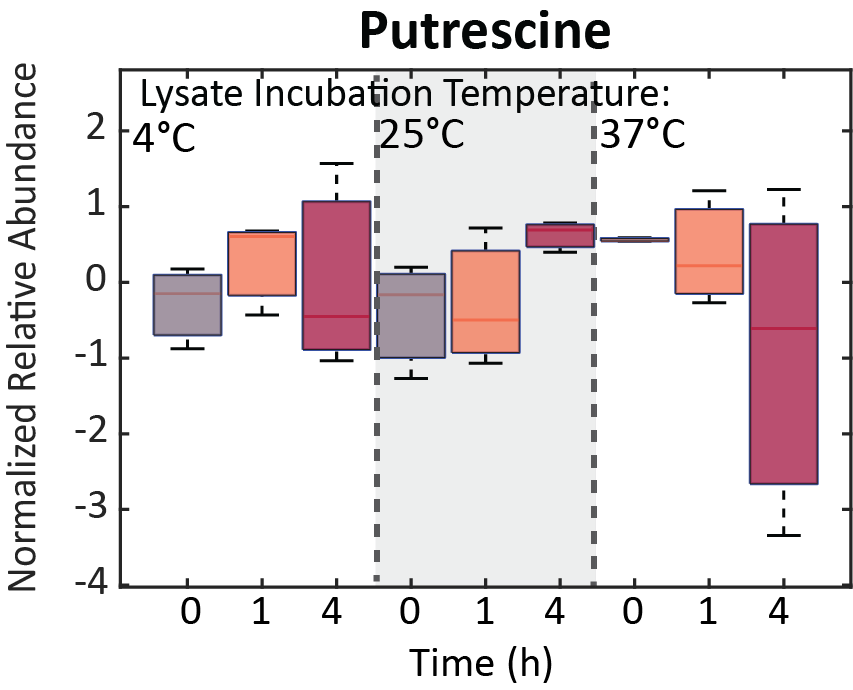

Figure S5. Relative abundances of putrescine in CFE reaction samples with lysates pre-incubated at different temperatures. Box and whisker plots depict the normalized peak areas, which are transformed using a generalized logarithm (base 2) and autoscaled. Red lines are the medians, boxes span the second and third quartiles of values. Error bars represent standard deviation of triplicate reactions.

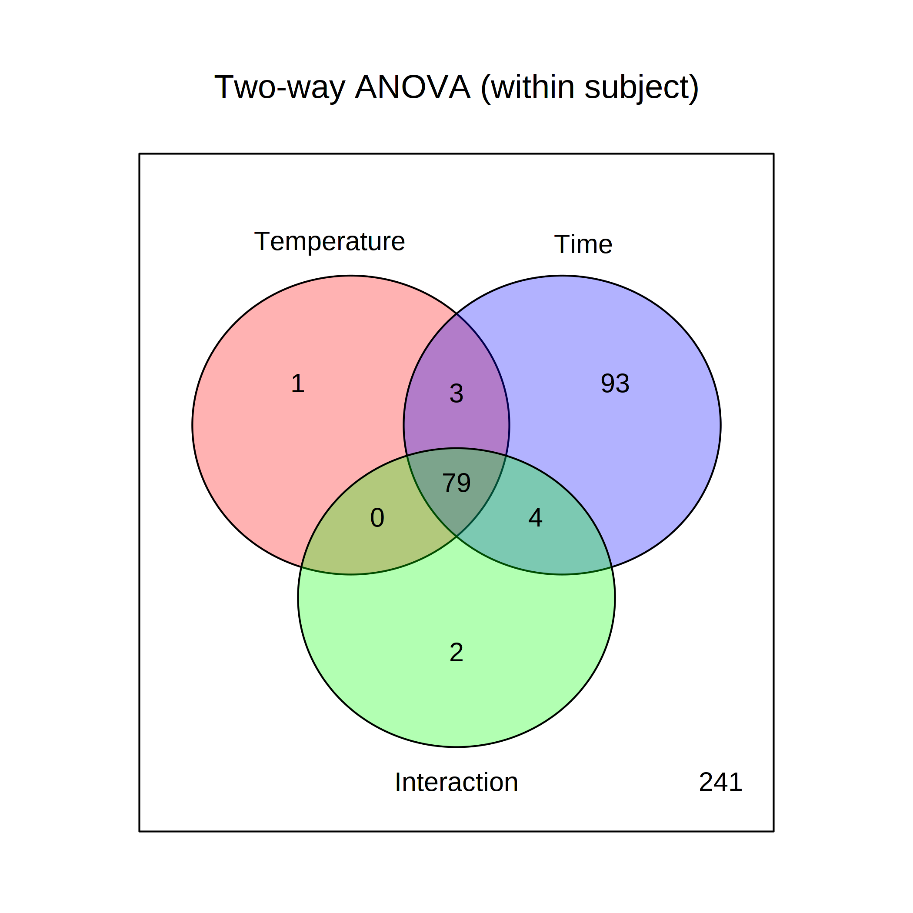

**A**

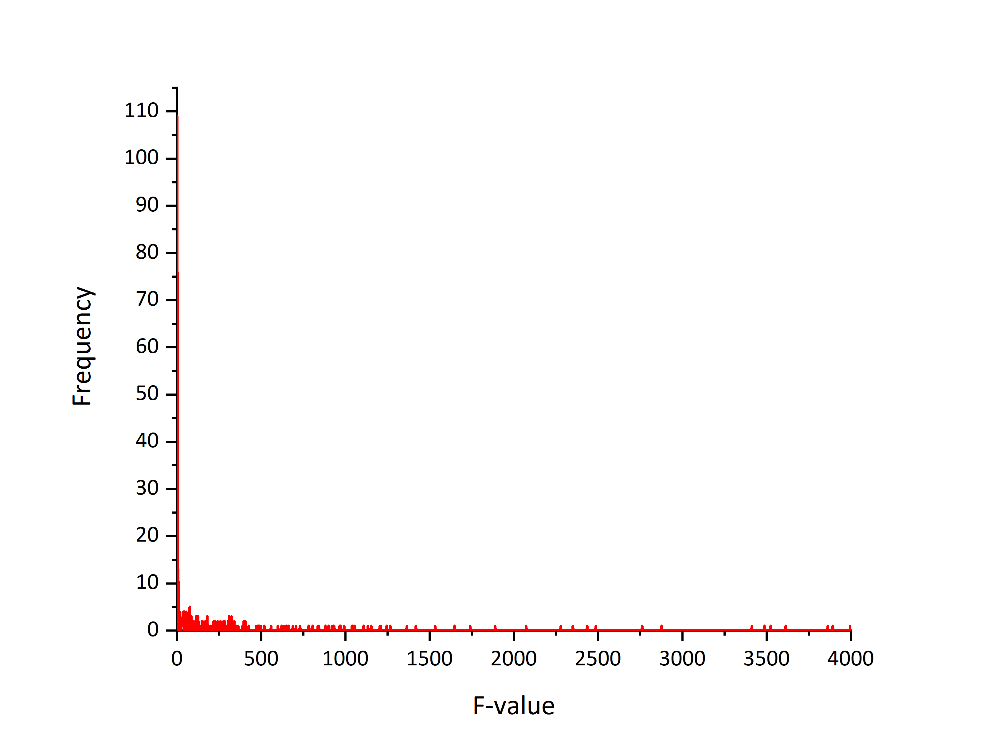

**B**

Figure S6. (A) Venn diagram reporting two-way ANOVA results and (B) the distribution of metabolite f-values from metabolomics data collected from CFE reactions run with lysates pre-incubated at 4, 25 and 37 °C for 6 hours. In (A), numbers represent counts of metabolites that have significant effects for each factor, assessed using FDR corrected *p*-values (<0.05). In (B), bin values of 1 were used to determine f-value frequency. The majority of f-values are between 0 and 9.

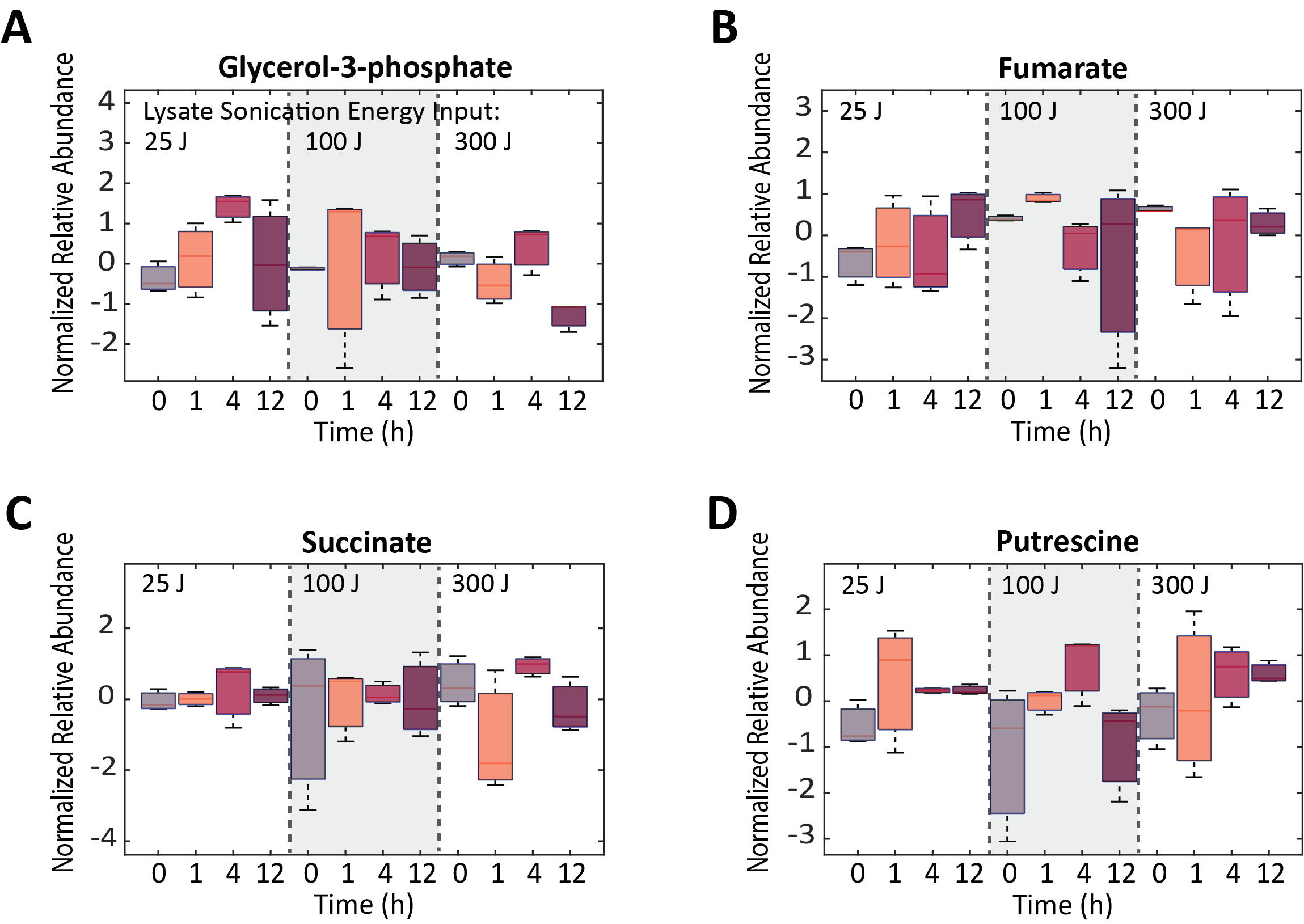

Figure S7. Relative abundances of (A) glycerol-3-phosphate, (B) fumarate, (C) succinate, and (D) putrescine in CFE reactions using differently sonicated lysates. Box and whisker plots depict the normalized peak areas, which were transformed using a generalized logarithm (base 2) and autoscaled. Red lines are the medians, boxes are the second and third quartile of values. Error bars represent standard deviation of triplicate reactions.

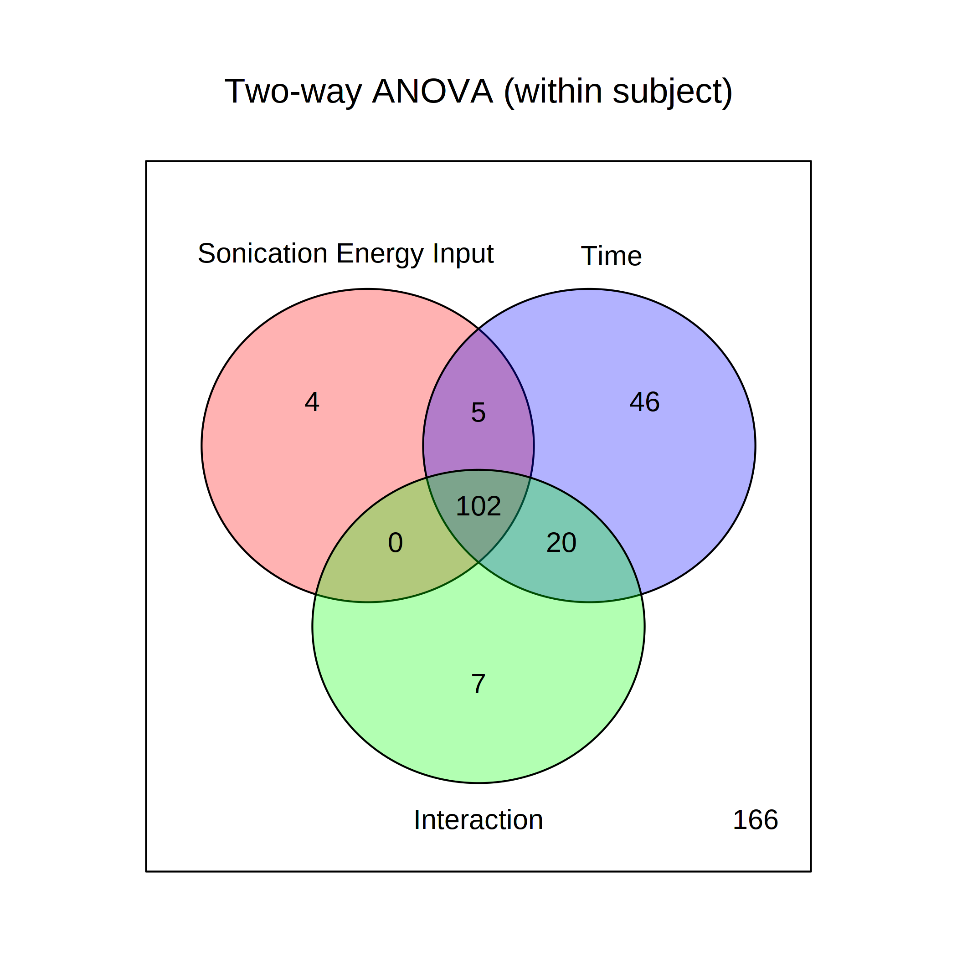

**A**

**B**

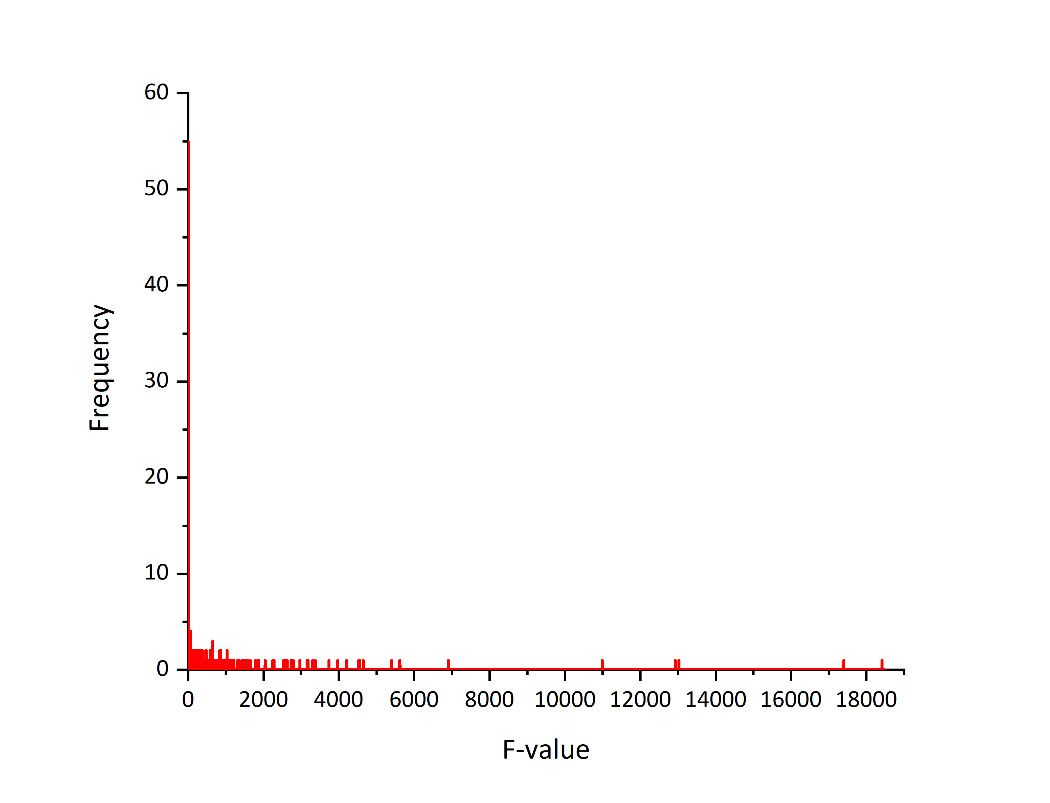

**Figure S8.** (A) Venn diagram reporting two-way ANOVA results and (B) the distribution of metabolite f-values from metabolomics data collected from CFE reactions run with lysates sonicated with different energy inputs. In (A), numbers represent counts of metabolites that have significant effects for each factor, assessed using FDR corrected *p*-values (<0.05). In (B), bin values of 1 were used to determine f-value frequency. The majority of f-values are between 0 and 15.

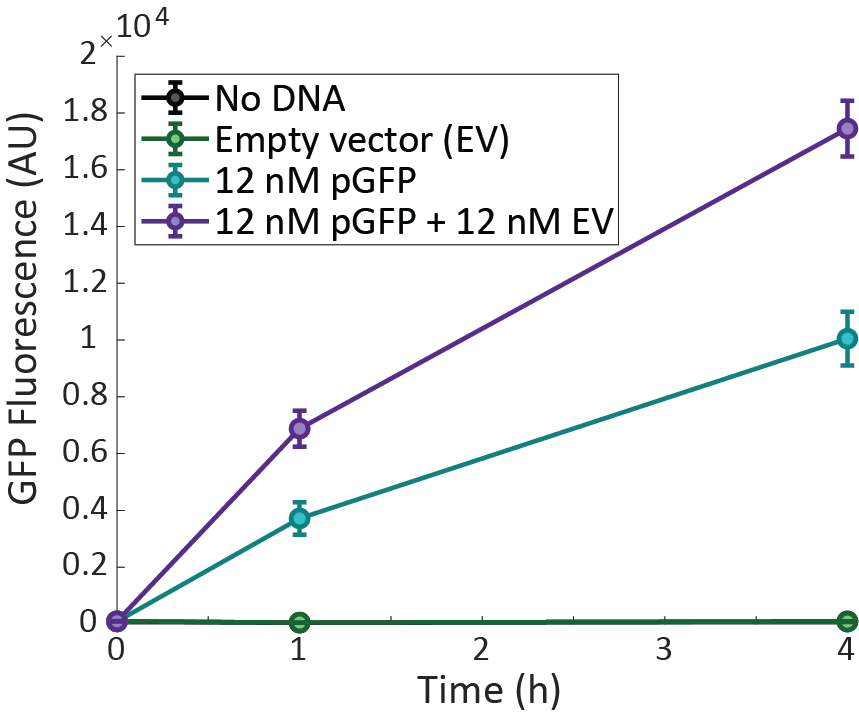

Figure S9. GFP production changes for reactions run with additional, non-reporter plasmids present. pGFP is pJL1s70, with expression controlled by a standard *E. coli* σ^70^ promoter. Error bars represent standard deviation of triplicate reactions.

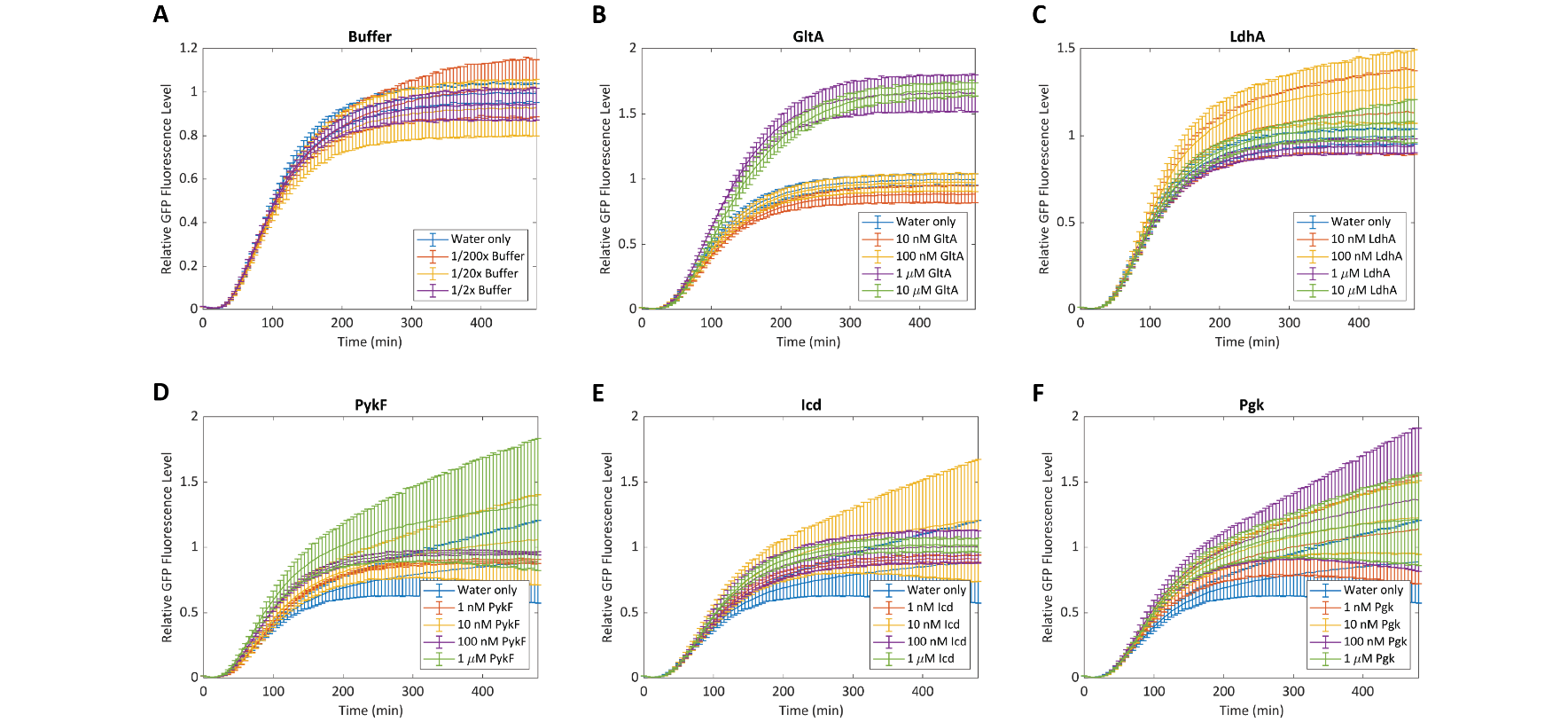

**Figure S10.** Optimization of enzyme supplementation levels in a CFE reaction. Different concentrations of (B) GltA, (C) LdhA, (D) PykF, (E) Icd, and (F) Pgk were added to CFE reactions and fluorescence was measured over 8 hours. Protein storage buffer at dilutions equivalent to those that would be added during enzyme supplementation were also tested (A) to control for potential protein expression improvements due to storage buffer components. 1 µM GltA and 100 nM LdhA statistically significantly improve endpoint GFP production. GFP expression was controlled by a standard *E. coli* σ^70^ promoter (pJL1s70). Error bars represent standard deviation of triplicate reactions.

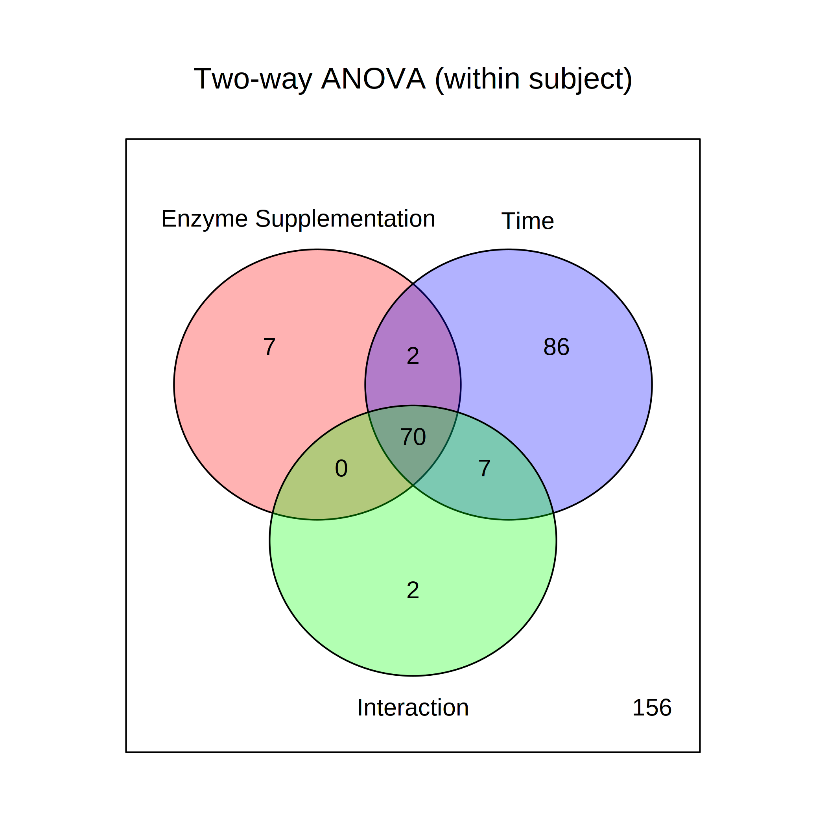

**A**

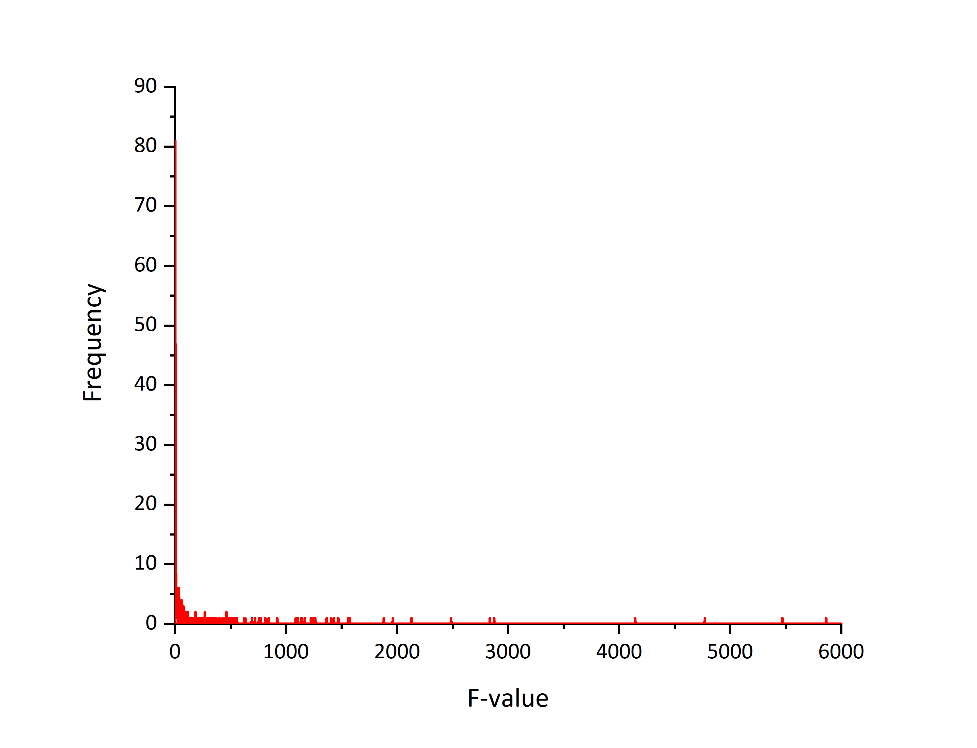

**B**

Figure S11. (A) Venn diagram reporting two-way ANOVA results and (B) the distribution of metabolite f-values from metabolomics data collected from CFE reactions run with supplemented enzymes. In (A), numbers represent counts of metabolites that have significant effects for each factor, assessed using FDR corrected *p*-values (<0.05). In (B), bin values of 1 were used to determine f-value frequency. The majority of f-values are between 0 and 7.
